# Whole Genomes Reveal Evolutionary Relationships and Mechanisms Underlying Gene-Tree Discordance in *Neodiprion* Sawflies

**DOI:** 10.1101/2023.01.05.522922

**Authors:** Danielle K. Herrig, Kim L. Vertacnik, Ryan D. Ridenbaugh, Kathryn M. Everson, Sheina B. Sim, Scott M. Geib, David W. Weisrock, Catherine R. Linnen

## Abstract

Rapidly evolving taxa are excellent models for understanding the mechanisms that give rise to biodiversity. However, developing an accurate historical framework for comparative analysis of such lineages remains a challenge due to ubiquitous incomplete lineage sorting and introgression. Here, we use a whole-genome alignment, multiple locus-sampling strategies, and locus-based and SNP-based species-tree methods to infer a species tree for eastern North American *Neodiprion* species, a clade of pine-feeding sawflies (Order: Hymenopteran; Family: Diprionidae). We recovered a well-supported species tree that—except for three uncertain relationships—is robust to different strategies for analyzing whole-genome data. Despite this consistency, underlying gene-tree discordance is high. To understand this discordance, we use multiple regression to model topological discordance as a function of several genomic features. We find that gene-tree discordance tends to be higher in regions of the genome that may be more prone to gene-tree estimation error, as indicated by a lower density of parsimony-informative sites, a higher density of genes, a higher average pairwise genetic distance, and gene trees with lower average bootstrap support. Also, contrary to the expectation that discordance via incomplete lineage sorting is reduced in low-recombination regions of the genome, we find a *negative* correlation between recombination rate and topological discordance. We offer potential explanations for this pattern and hypothesize that it may be unique to lineages that have diverged with gene flow. Our analysis also reveals an unexpected discordance hotspot on Chromosome 1, which contains several genes potentially involved in mitochondrial-nuclear interactions and produces a gene-tree that resembles a highly discordant mitochondrial tree. Based on these observations, we hypothesize that our genome-wide scan for topological discordance has identified a nuclear locus involved in a mito-nuclear incompatibility. Together, these results demonstrate how phylogenomic analysis coupled with high-quality, annotated genomes can generate novel hypotheses about the mechanisms that drive divergence and produce variable genealogical histories across genomes.

---

It has long been recognized that gene trees need not match species trees due to stochastic sorting of ancestral polymorphism (i.e., incomplete lineage sorting), hybridization and introgression, horizontal gene transfer, and gene duplication and loss (Avise et al. 1987; Maddison 1997). An important implication of gene tree-species tree discordance is that data from many unlinked loci are necessary to accurately reconstruct divergence histories, but large molecular datasets were prohibitively expensive to generate using pre-genomic methodologies. Together, next-generation sequencing and the multi-species coalescent model have revolutionized molecular phylogenetics (Edwards 2009; McCormack et al. 2013; Edwards et al. 2016; Rannala et al. 2020). Next-generation sequencing made it possible to comprehensively sample the “cloud of gene histories” (Maddison 1997) embedded within a species tree. As molecular datasets grew, however, debates emerged over the best way to handle large multilocus datasets with heterogeneous phylogenetic signal (Gatesy and Springer 2013; Edwards et al. 2016). One approach that emerged is to concatenate all loci into a single supermatrix to be analyzed using standard phylogenetic methods. Concatenation is computationally efficient and can produce an accurate species tree when incomplete lineage sorting is minimal (Edwards et al. 2007; Kubatko and Degnan 2007; Leaché and Rannala 2011). However, under higher levels of incomplete lineage sorting, concatenation is statistically inconsistent and can produce a highly supported, but incorrect, species tree topology (Roch and Steel 2015). By modeling the coalescent process that gives rise to heterogeneous gene trees, the multi-species coalescent model (Rannala and Yang 2003) yields more accurate estimates of species-tree topologies and divergence times (Ogilvie et al. 2017; Jiang et al. 2020; Rannala et al. 2020).

Despite much progress, a central challenge in coalescent-based species-tree analysis is that no model accounts for all possible sources of gene-tree discordance. So-called “full-likelihood methods” (e.g., BEST and *BEAST; Liu 2008; Heled and Drummond 2010) accommodate both uncertainty in gene-tree estimation and incomplete lineage sorting by simultaneously modelling nucleotide substitution and the coalescent process, but these approaches remain computationally burdensome (Liu et al. 2015; Rannala et al. 2020). Moreover, available nucleotide substitution models may be inadequate for describing complex patterns of sequence evolution, particularly for loci evolving under strong selection (Whelan and Goldman 2001; Yang and Nielsen 2002; Whelan 2008; Chi and Liberles 2016; Echave et al. 2016; Reddy et al. 2017; Dornburg et al. 2019; Karin et al. 2020). In lieu of full-likelihood methods, many researchers use approximate approaches based on the multi-species coalescent model (e.g., MP-EST and ASTRAL, Liu et al. 2010; Zhang et al. 2018). These methods use estimated gene trees for species-tree inference and assume that input gene trees are accurate and that there is no recombination within, but free recombination between loci. These assumptions create a trade-off for sampling loci: shorter loci are more likely to satisfy the “no intralocus recombination” assumption, but less likely to yield sufficient information for accurate gene-tree inference (Chou et al. 2015).

Another source of gene-tree discordance that is widespread in nature is introgression (Harrison and Larson 2014; Leaché et al. 2014; Fontaine et al. 2015; Mallet et al. 2016; Edelman et al. 2019; Hibbins and Hahn 2022). The multi-species coalescent model has been extended to include interspecific gene flow (Hey and Nielsen 2004; Yu et al. 2014), but full-likelihood implementations of these models (Jones 2018; Wen and Nakhleh 2018; Wen et al. 2018; Zhang et al. 2018a; Flouri et al. 2020) are computationally demanding (Flouri et al. 2020; Hibbins and Hahn 2022). For this reason, heuristic approaches for detecting introgression (e.g., SNaQ, ABBA-BABA tests, or HyDe; Green et al. 2010; Durand et al. 2011; Solís-Lemus et al. 2016; Blischak et al. 2018) are often used in conjunction with other species-tree methods that assume discordance is due to incomplete lineages sorting (e.g., Edelman et al. 2019; Meleshko et al. 2021). Although no species-tree method accounts for all sources of gene-tree discordance, coalescent-based methods nevertheless appear to perform reasonably well even when model assumptions are violated (Lanier and Knowles 2012; Chou et al. 2015; Adams et al. 2018; Long and Kubatko 2018; Borges et al. 2022; Yan et al. 2022).

In addition to creating new analytical challenges, genome-scale datasets also reveal that gene-tree discordance is unevenly distributed across genomes (Pollard et al. 2006; White et al. 2009; Fontaine et al. 2015; Edelman et al. 2019; Small et al. 2020). One potential source of heterogeneous discordance is gene-tree estimation error, which may vary across the genome due to local base composition, repetitive sequence content, and density of phylogenetically informative sites (Betancur-R. et al. 2013). The genomic landscape of discordance is also likely to be influenced by the interplay between natural selection, gene flow, and recombination. For example, convergent molecular evolution (Christin et al. 2007; Castoe et al. 2009) and adaptive introgression (Pardo-Diaz et al. 2012; Jones et al. 2018; Oziolor et al. 2019) can increase gene-tree discordance at selected and linked sites. Conversely, selection in ancestral lineages may reduce the potential for discordance via incomplete lineage sorting because selection on a locus reduces the effective population size (*N*_*e*_) at linked neutral sites (Smith and Haigh 1974; Charlesworth et al. 1993; Slatkin and Pollack 2006; Stukenbrock et al. 2011; Pease and Hahn 2013; Dutheil et al. 2015). For this reason, gene-tree discordance via incomplete lineage sorting may be reduced in low-recombination regions of the genome (Pease and Hahn 2013). Gene-tree discordance via introgression may also be reduced in low-recombination regions of the genome. This is because neutral or positively selected variants in low-recombination regions of the genome are more likely to be linked to deleterious alleles that prevent introgression (Nachman and Payseur 2012; Brandvain et al. 2014; Schumer et al. 2018; Li et al. 2019; Martin et al. 2019). Overall, research to date suggests that gene-tree discordance should correlate with genomic variables—such as gene density, base composition, recombination rate, and the number of informative sites—that influence the probability of gene-tree estimation error, incomplete lineage sorting, and introgression. However, the relative contribution of different genomic predictor variables to genome-wide patterns of gene-tree discordance remains an open question.

Here, we revisit a classic case study in messy species-tree inference (Linnen and Farrell 2007, 2008b, 2008a; Linnen 2010) armed with a high-quality reference genome, whole-genome resequencing data for 19 species, and newer species-tree methods. Specifically, we focus on the eastern North American “*Lecontei*” clade of *Neodiprion* sawflies (Order: Hymenoptera, Family: Diprionidae). Previous studies suggest that this clade radiated with substantial gene flow sometime within the last 2-10 million years (Linnen and Farrell 2007, 2008b; Bendall et al. 2022). Upon colonizing eastern North America, population divergence and speciation were likely driven by rapid adaptation to new *Pinus* hosts (Linnen and Farrell 2010; Bagley et al. 2017; Bendall et al. 2017; Glover et al. 2023). In addition to variation in host use, there is intra- and interspecific variation in larval and adult morphology, behavior, and overwintering strategy (Coppel and Benjamin 1965; Knerer and Atwood 1973). This phenotypic variation, coupled with a rich natural history literature and experimentally tractable species that can be reared and crossed in the lab (Knerer 1984; Bendall et al. 2017; Linnen et al. 2018), makes *Neodiprion* an excellent model for characterizing both the genetic mechanisms and evolutionary drivers of population differentiation and speciation. For robust inferences about evolution in this group, an accurate species-tree estimate is essential.

The first informal phylogenetic hypothesis for *Neodiprion* consisted of five named species complexes (*lecontei, pinusrigidae, pratti, abbotii*, and *virginianus*) based on shared morphological and ecological traits (Ross 1955). Over fifty years later, these proposed species groups were evaluated with DNA sequence data from one mitochondrial locus and three nuclear genes. As expected under a scenario of rapid and recent divergence with substantial gene flow, gene-tree topologies differ among the four loci (Linnen and Farrell 2007, 2008a). However, gene-tree discordance is especially pronounced between the mitochondrial locus and the three nuclear loci, likely due to extensive mitochondrial introgression (Linnen and Farrell 2007). To obtain a species-tree estimate from the remaining three nuclear loci, Linnen and Farrell (2008a) used multiple species-tree methods (Takahata 1989; Maddison 1997; Maddison and Knowles 2006; Edwards et al. 2007; Liu and Pearl 2007). Overall, these analyses yielded consistent support for two of Ross’s proposed species groups (*lecontei* and *pinusrigidae*), mixed support for the *virginianus* and *pratti* species groups, and no support for the *abbotii* species group. Although different methods and subsets of individuals per species produced different topologies, Linnen and Farrell (2008a) used areas of agreement across phylogenetic analyses to propose a provisional species tree for the *Lecontei* group (Fig. 1).

**Figure 1:**
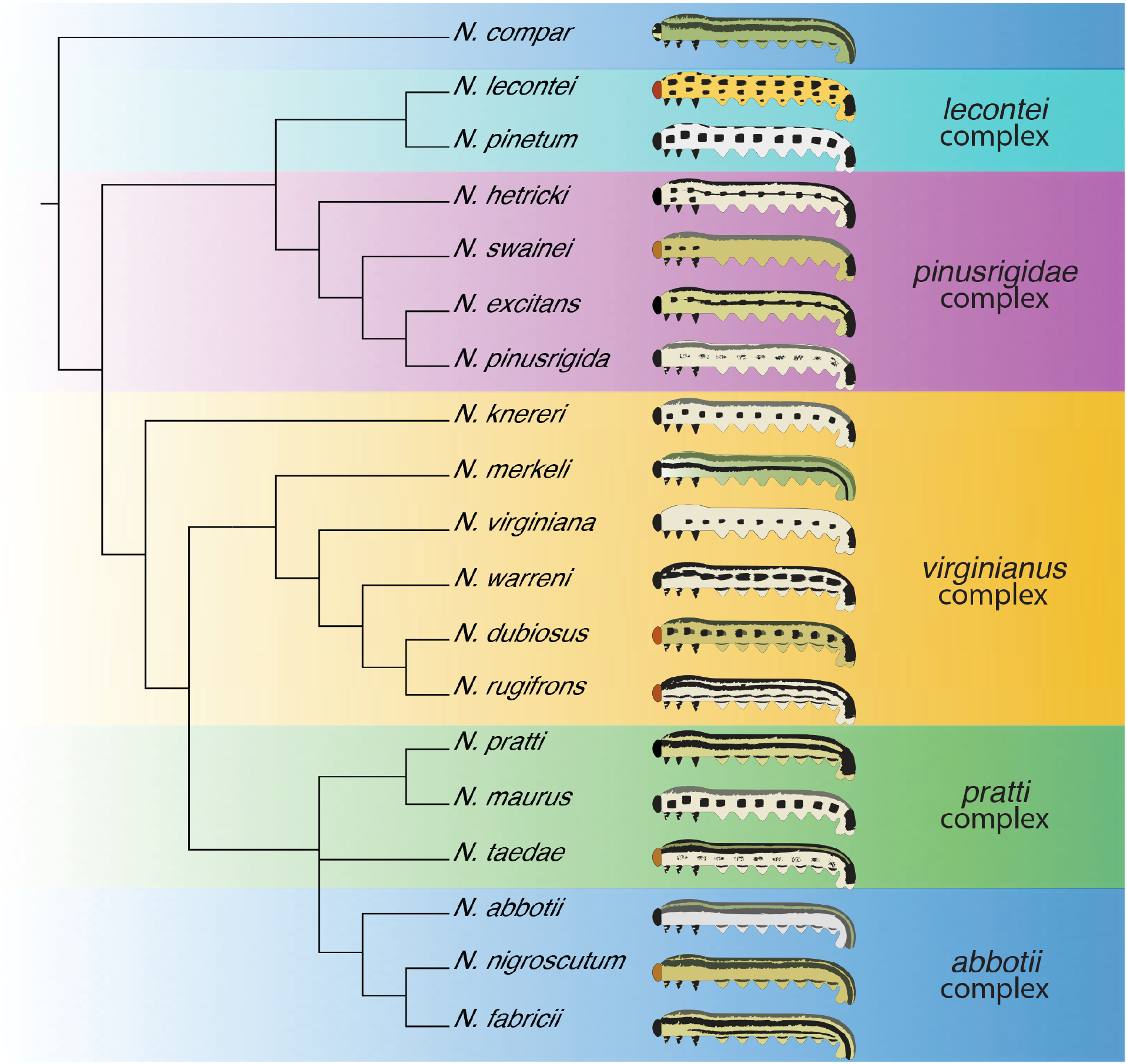
Proposed *Neodiprion* species complexes and species tree estimated from three nuclear loci. Colors correspond to proposed species complexes (Ross 1955) and tree topology corresponds to proposed topology in Linnen and Farrell 2008a. Cartoons depict typical larval color pattern for each species.

In this study, we expand the four-locus *Lecontei-*group dataset to the entire genome. Our study has three goals. First, we take advantage of a small, tractable genome (∼272 Mb) to evaluate how inferred species-tree topologies are influenced by: (1) different strategies for parsing large contiguous chromosomes into separate loci (from 5 kb to 1 Mb loci) for use in ASTRAL-III analyses; (2) different analysis methods; specifically, locus-based (ASTRAL-III) vs. SNP-based (SVDquartets) approaches; and (3) subsampling the genome in ways that mimic reduced-representation approaches such as exon capture (approximated by sampling coding exons only) and restriction-associated DNA sequencing (approximated by subsampling SNPs). Second, taking advantage of a new reference-quality assembly we generated to facilitate this work, we investigate genome-wide patterns of gene-tree discordance. We use this approach to evaluate genomic correlates of discordance and identify gene-tree “outliers” that may point to hotspots for adaptation from standing variation, adaptive introgression, or genetic incompatibilities. Third, we integrate all species-tree results to suggest an updated species tree for comparative work, highlighting areas of remaining uncertainty that are sensitive to sampling strategy and species-tree methodology. Although we identify multiple genomic correlates of topological discordance and a pronounced discordance hotspot on one chromosome, we also find that outside of a few uncertain relationships, sampling and analysis strategy have a minimal impact on the inferred species-tree topology. Based on these findings, we highlight priorities for future work on this system and make recommendations for other genome-wide phylogenomic analyses.

## Materials & Methods

### *Assembly and Annotation of a Reference-Quality Genome for* Neodiprion lecontei

#### DNA extraction and library preparation

We obtained samples for DNA extraction from haploid male siblings that were the progeny of a single virgin *N. lecontei* female that had been collected in Lexington, KY (38°00’50.4”N 84°30’14.4”W) as a larva and lab-reared to adulthood. To maximize sawfly DNA yields and minimize host plant material in the gut, we flash-froze male larvae that were either in the final, non-feeding instar or dissected from freshly spun cocoons. We extracted genomic DNA from a single haploid male with a MagAttract HMW DNA Kit (Qiagen, Hilden Germany) using the fresh or frozen tissue protocol. To further improve sample purity, we performed a 2.0x bead clean-up using polyethylene glycol containing solid-phase reversible immobilization beads solution for each sample (DeAngelis et al. 1995). We quantified double-stranded DNA using a dsDNA Broad Range (BR) Qubit assay and assessed using the fluorometer feature of a DS-11 Spectrophotometer and Fluorometer (DeNovix Inc, Wilmington, DE, USA). We quantified DNA purity using the UV-Vis spectrometer feature on the DS-11, which reported OD 230/260/280 ratios.

We sheared DNA to a mean size distribution of ∼20 kb using a Diagenode Megaruptor 2 according to the manufacturer’s protocol, (Denville, New Jersey, USA) and sized sheared DNA on a Fragment Analyzer (Agilent Technologies, Santa Clara, California, USA) using the High Sensitivity (HS) Large Fragment kit. Sheared DNA was the starting input for PacBio SMRTBell library preparation using the SMRTbell Express Template Prep Kit 2.0 according to the manufacturer’s protocol (Pacific Biosciences, Menlo Park, California, USA). The final library was shipped to the USDA-ARS Genomics and Bioinformatics Research Unit in Stoneville, Mississippi, USA where it was sequenced on one Pacific Biosciences 8M SMRT Cell on a Sequel II system (Pacific Biosciences, Menlo Park, California, USA) with a pre-extension time of 2 hours and a movie collection time of 30 hours.

In a parallel pipeline, we prepared an enriched chromosome conformation capture (HiC) library using another *N. lecontei* sample from the same haploid male family. Briefly, we crosslinked tissue using the Arima HiC low input protocol and performed proximity ligation using the Arima HiC Kit (Arima Genomics, San Diego, California, USA). After proximity ligation, we sheared the DNA using a Diagenode Bioruptor and then size-selected to enrich for DNA fragments of 200-600bp. We prepared an Illumina library from the sheared and size-selected DNA using the Swift Accel NGS 2S Plus kit (Integrated DNA Technologies, Coralville, Iowa, USA). The final Illumina HiC library was sequenced on a NovaSeq 6000 at the Hudson Alpha Genome Sequencing Center (Huntsville, Alabama, USA) with paired-end 150 bp (PE150) reads. We trimmed sequence reads for Illumina adapter artifacts using the Illumina BaseSpace software (Illumina, San Diego, California, USA).

#### Genome assembly

Following sequencing, circular consensus sequence (CCS) calling was performed on the raw subreads generated by the Sequel II system using the SMRTLink v8.0 software (Pacific Biosciences, Menlo Park, California, USA). We filtered the resulting CCS reads for adapter contamination using the software HiFiAdapterFilt v2.0 (Sim et al. 2022). The filtered dataset served as input sequences for the HiFi data assembly software HiFiASM v0.16.1-r375 (Cheng et al. 2021) using default parameters. We converted the resulting contig assembly (.gfa format) to .fasta format using the software any2fasta (Seeman, 2018 https://github.com/tseemann/any2fasta).

To produce the HiC scaffolded assembly, we mapped paired Illumina HiC reads to the HiFi derived contig assembly using the mem function of the software BWA. We removed PCR duplicate artifacts from the resulting .sam file using the software samblaster (Faust and Hall 2014). We then used the resulting .bam file as the input file to the Phase Genomics Matlock suite of HiC functions (Kronenberg and Sullivan, 2018 https://github.com/phasegenomics/matlock) (Phase Genomics, Seattle, Washington, USA), which converted mapped reads into a HiC format that could be converted to a .hic and .assembly file using the 3d-dna script ‘run-assembly-visualizer.sh’ and visualized using the software Juicebox v1.11.08 (Durand et al. 2016). Manual edits to the HiC scaffold assembly were performed using Juicebox v1.11.08 and changes were applied to the assembly using the Phase Genomics script ‘juicebox_assembly_converter.py’ (https://github.com/phasegenomics/juicebox_scripts).

#### Assembly quality analysis

To estimate genome size, we performed K-mer distribution analysis on the raw data using the k-mer counting software KMC (Deorowicz et al. 2013) and GenomeScope v.2.0 (Ranallo-Benavidez et al. 2020). To evaluate the assembly for duplicate contigs, we performed k-mer spectra analysis with the K-mer Analysis Toolkit (KAT) (Mapleson et al. 2016) using the raw data and the contig assembly. We evaluated duplicate content and genome completeness with BUSCO v5.0 in ‘genome’ mode for the Metazoa, Arthropoda, Insecta, Endopterygota, and Hymenoptera ortholog sets (Manni et al. 2021). To characterize the read depth of each contig, we used the mapping software minimap2 (Li 2018) with the contig assembly and raw HiFi CCS read set. We used the NCBI nucleotide (nt) database (accessed 2017-0605) and the UniProt Reference Proteomes database (accessed 2020-03) to assign each contig to a taxonomic class using BLAST+ (Camacho et al. 2009) in blastn mode and Diamond (Buchfink et al. 2021) in blastx mode, respectively. Results from the hierarchical BUSCO analysis, read depth analysis, and taxon assignment were visualized using BlobTools2 (Laetsch and Blaxter 2017), and summarized using the python script blobblurb (https://github.com/sheinasim/blobblurb).

#### Gene annotation

The *N. lecontei* genome was submitted to NCBI for annotation using previously deposited RNAseq data (Herrig et al. 2021) for annotations. The NCBI *Neodiprion lecontei* Annotation Release 101 was completed using the NCBI Eukaryotic Genome Annotation Pipeline Software version 9.0. Briefly, BUSCO v4.1.4 was run in protein mode on the annotated gene set and the longest gene was retained. WindowMasker (Morgulis et al. 2006)was used to mask the genome (26.24% masked). Previously deposited transcriptome sequences from *Neodiprion* (including 77 specific tissues isolated from *N. lecontei* at five life stages), RefSeq proteins, and GenBank Insecta proteins were then aligned to the masked genome using Splign, minimap2, or ProSplign (Kapustin et al. 2008; Li 2018). The alignments were then passed to Gnomon (Souvorov et al. 2010) for gene predictions. 14,732 genes and pseudogenes were identified in this process including 11,969 protein-coding genes.

### Taxonomic Sampling, Library Preparation and Sequencing, and Reference-Anchored Alignment

We extracted fresh DNA from ethanol-preserved exemplars from 19 *Neodiprion* species, often from the same individuals or colonies as used in earlier phylogenetic studies of this genus (Linnen and Farrell 2007). Larval individuals had been collected in the United States and Canada (Table S1) and stored in ethanol at -20ºC. Sampling included all species in the eastern North American “*Lecontei*” clade except *N. insularis* and *N. cubensis*, both endemic to Cuba. We included the western North American species *N. autumnalis* an outgroup. Based on the presence of heterozygous sites in Sanger-sequence data from three nuclear loci, all individuals included in this study were diploid.

We dissected tissue from the prolegs and the ventral region of the larvae, avoiding the gut region. We then ground liquid nitrogen-frozen tissue with pestles made from 1 mL micropipette tips, and incubated the resulting powder in CTAB buffer with proteinase K and RNase A. We extracted DNA using phenol-chloroform-isoamyl alcohol, dried the ethanol precipitate overnight, and resuspended in TE buffer. We assessed DNA integrity with a 0.7% agarose gel, DNA purity with the 260/280 ratio, and DNA concentration with a Quant-iT dsDNA High-Sensitivity fluorescence assay (Thermo Fisher Scientific). The Georgia Genomics and Bioinformatics Core (Athens, GA, USA) prepared and sequenced one small-insert DNA library for each species. Libraries had a mean fragment size of 619 bp and were sequenced on Illumina NextSeq 500 with PE150 reads. Sequencing produced 14-27 million filtered reads per individual.

To obtain a multi-genome alignment, we used a pseudo-reference-based approach, with our annotated, reference quality *N. lecontei* genome (described above) serving as the reference. This approach is appropriate for this clade because synteny is high (unpublished data) and genetic divergence among species is low (see results). Our pipeline was based on the published Pseudo-It pipeline (Sarver et al. 2017) (Fig. S1) with some modifications. Briefly, we first used bowtie2 v2.4.1 (Langmead and Salzberg 2012) to map reads from each species to the *N. lecontei* reference genome. To allow for divergence between reads and the *N. lecontei* reference, we initially allowed a mismatch in the seed and “local” mapping options in bowtie2. New variants (excluding indels) were incorporated using samtools v1.10 (Li et al. 2009) and bcftools v1.10.2 (Li 2011). In a second round of mapping, this process was repeated using the first iteration of the genome for each species as the new reference genome. The third round of mapping removed the seed mismatch. The fourth and fifth iterations required end-to-end mapping. After the fifth iteration, we replaced any nucleotide that had a read depth less than 4 or that had excessively high mapping depth (highest 1% of depths for each species) with an “N” using a custom script. Unless otherwise noted, all bioinformatics commands and scripts can be found on the LinnenLab GitHub page under the Herrig_etal_NeodiprionPhylogeny repository.

### Dataset Preparation and Phylogenetic Inference

Our genome assembly and pseudo-reference approach produced 20 aligned genomes: the *de novo N. lecontei* genome and 19 pseudo-reference genomes (18 *Lecontei* clade species plus outgroup *N. autumnalis)*. To explore how genomic sampling strategy and analysis method affects species-tree inference, we sampled the aligned genomes in three ways. First, we used bedtools v2.30.0 (Quinlan 2014) to divide the seven *Neodiprion* chromosomes into non-overlapping windows of different sizes: 5 kb, 10 kb, 50 kb, 100 kb, 500 kb, and 1 Mb. We then used IQ-TREE v2.1.2 (Minh et al. 2020) to estimate a maximum-likelihood gene tree for each window, allowing IQ-tree to select the best-fit substitution model for each window (-m MFP option). For each tree, we calculated support using 1,000 bootstraps using the ultrafast bootstrap approximation. Because some regions of the genome had high levels of missing data, we used custom python scripts to exclude gene trees for which the average amount of missing data (Ns) per species was more than 10%. We then input the remaining trees into ASTRAL-III v5.7.3 (Zhang et al. 2018b) to produce a single coalescent-based species tree. We obtained quartet support for each node using the –t 1 option.

Second, to approximate a dataset of protein-coding genes analogous to an RNAseq or exon-capture phylogenomic dataset, we used gffread v0.11.7 with the –w flag to write fasta files with spliced exons for each transcript for each species using the NCBI *Neodiprion lecontei* Annotation Release (iyNeoLeco1.1 RefSeq GCF_021901455.1). Custom scripts were used to define and keep the isoform with the most parsimony informative sites. We again used IQ-TREE to estimate a maximum-likelihood gene tree and bootstrap support for each gene, excluding genes containing fewer than 10 parsimoniously informative sites. As an additional step to filter out uninformative loci, we filtered gene trees based on average bootstrap support across the tree, retaining only those trees that had average bootstrap support values ≥ 60%, 70%, or 80%. Finally, we used ASTRAL-III to estimate a species tree and quartet support for each node for each bootstrap value cutoff.

Third, we called single nucleotide polymorphisms (SNPs) across the entire genome using SNP-sites v2.5.1 (Page et al. 2016). We then filtered the data to exclude SNPs that were absent in more than 10% of species and sites with more than two alleles. In addition to analyzing all SNPs (which likely contains tightly linked sites), we produced additional datasets with one SNP sampled every 1 kb, 5 kb, 10 kb, 50 kb, or 100 kb using SNP-sites, with more sparsely sampled SNPs on par with a dataset that might be generated via RADseq. We transformed each of the six datasets into nexus format and used SVDquartets (Chifman and Kubatko 2014), implemented in PAUP v4.0a (Swofford 2000), to produce a species tree for each dataset allowing it to evaluate all possible quartets, with support determined via 10,000 bootstraps. Ambiguous sites were handled using the “Distribute” option.

### Visualization of Gene-Tree Heterogeneity

We visualized variation in gene-tree topology across the genome in two ways. First, to visualize overall topological discordance among gene-trees produced by 500 kb windows, we overlaid each gene tree over one another using the program densiTree (part of the phangorn package, v2.7.0) (Bouckaert 2010). To prepare these trees for visualization, ape v5.5 (Paradis and Schliep 2019), phangorn v2.7.0 (Schliep 2011), and phytools v0.7-90 (Revell 2012) packages built in R v3.6.2 were used to input gene trees, root them using *N. autumnalis*, extract trees as phylo objects, and standardize branch lengths (using the “Grafen” method in compute.brlen). To evaluate the potential for introgression to contribute to discordant topologies revealed in the densiTree plot, we used the DtriosCombine utility in the Dsuite v0.4 package (Malinsky et al. 2021) to calculate genome-wide D statistics (ABBA-BABA) for all possible trios of species, with the 500 kb ASTRAL-III species-tree as the guide tree. To obtain a single D-statistic estimate for each species pair, we averaged genome-wide D-statistics from all possible trios for that species pair that were given by Dsuite. We then created a heatmap of pairwise D-statistics for comparison to the densiTree plot and to published estimates of pairwise mitochondrial introgression rates (population migration rate, *2Nm*) obtained via the program IM (Hey and Nielsen 2004; Linnen and Farrell 2007).

Second, to visualize how topological discordance between gene trees and the inferred species tree varies across the genome, we calculated Robinson-Foulds distances (Robinson and Foulds 1981) between the gene-tree estimated for each window and the overall species tree estimated using all window-based trees using the treedist function in R package Phangorn v2.8.1. We then used the R package chromPlot v3.6.2 (Oróstica and Verdugo 2016) to paint the chromosomes based on estimated Robinson-Foulds distances. To determine how genomic patterns of discordance varied as a function of window size, we computed distances and painted chromosomes for 1 Mb, 500 kb, 100 kb, and 50 kb windows.

### Genomic Correlates of Gene-Tree Discordance

To assess the potential for gene-tree estimation errors to contribute to topological discordance, we used custom scripts to compute several summary statistics for each window that reflect information content of alignments and confidence in gene-tree estimates. These statistics, pulled from IQ-tree output, include: the number of parsimoniously informative sites per window, the average bootstrap support across all nodes in the corresponding gene tree, and the proportion of nucleotides that were G or C (GC content). Variation in discordance can also be generated by heterogeneous selection, recombination rates, and introgression across the genome. To capture some of these potential sources of variation in discordance across the genome, for each 500 kb window we calculated: gene density, average pairwise genetic distance (K80), average D-statistics, and local recombination rate. To calculate gene density, we used the NCBI *Neodiprion lecontei* Annotation Release 1.1 (iyNeoLeco1.1 RefSeq GCF_021901455.1) to define the number of genes with start codons within each window using custom scripts. To calculate genetic distance, we used the dist.dna command within the R v3.6.2 package ape v5.5 (Paradis and Schliep 2019) and default parameters to calculate Kimura’s two-parameter (K80) distance (Kimura 1980) between every pair of species. To obtain a single genetic distance measure for each window, we then averaged all pairwise distances. To calculate a single D-statistic for each window, we ran a separate Dtrios analysis for each window, then averaged the resulting D-statistics across all trios available for each window.

To estimate local recombination rate, we used data from a mapping population that was previously used to identify quantitative trait loci for larval color traits that differ between two populations of *N. lecontei* (Linnen et al. 2018). The original analysis consisted of 503 SNPs genotyped in 429 F_*2*_ haploid males (like all hymenopterans, *Neodiprion* are haplodiploid) generated via double-digest restriction-associated DNA sequencing (Peterson et al. 2012). To increase marker density, we first mapped the raw sequencing reads (NCBI BioProject PRJNA434591, SRR6749156) to the new *N. lecontei* genome assembly (iyNeoLeco1.1). To identify fixed differences between the two parental populations, we first mapped reads from the cross parents (4 males; 4 females) and 10 F_1_ females to the reference genome using BWA-MEM v0.7.17 (Li 2013) with the –M option, followed by samtools v1.10 (Li et al. 2009) to convert the sam output to a bam file. Single nucleotide polymorphisms (SNPs) were called using mpileup in bcftools v1.9 (Li et al. 2009). We then used vcftools v0.1.16 (Danecek et al. 2011) to remove sites with indels and sites that were missing genotype information for more than 50% of the parents. To retain only putative fixed differences between the parent populations, we used vcftools to calculate the F_ST_ for each site and a custom python script (filterFst_sites.py) to retain sites with F_ST_ = 1, resulting in 21,887 SNPs. To further validate that these SNPs were fixed between populations, we required that at least one of the F_1_ females was heterozygous and that none of the F_1_ females were homozygous at a read depth of 10. We used a custom python script (checkF1hetstatus-readdepthcutoffforhomozygous.py) to drop any sites that did not meet these criteria, reducing our dataset to 13,462 SNPs. We next mapped F_2_ haploid male reads to the reference genome and called SNPs using BWA, samtools, and bcftools. For genotyping, we required a minimum read depth of 4 at each site and removed sites that had more than 50% missing information and that did not pass filtering in parents and F_1_ females using custom scripts (assignF2males_WY-singlemaleinput-minrd.py). In total, we retained 3,104 ancestry-informative SNPs that were called in at least 50% of F_2_ males.

We used R/qtl (Broman et al. 2003) to remove sites with identical genotypes to another site, sites exhibiting segregation distortion, and individuals that were missing genotypes at more than 50% of markers, producing a final dataset of 1436 markers genotyped in 402 F_2_ males. Marker location was inferred based on physical location in the reference genome. To estimate a genetic map with positions of markers in centiMorgans (cM), we used the “quickEst” function in ASMap v1.0-4 (Taylor and Butler 2017), with a Kosambi mapping function. To estimate local recombination rate, we used MareyMap Online (Rezvoy et al. 2007; Siberchicot et al. 2017) with the sliding window option. We then used the recombination rate estimate for the midpoint of each window for our regression analysis.

To determine which genomic summary statistics predicted topological discordance, we used the MASS v7.3-54 package in R v3.6.2 to fit a negative binomial model to the topological discordance data (Robinson-Foulds distances), with all seven summary statistics as predictor variables. We chose a negative binomial distribution because (1) Robinson-Foulds distances are counts (number of partitions that are not shared between the trees), and (2) initial model fitting indicated that a Poisson model was not appropriate for the data (the residual deviance divided by the degrees of freedom was > 1 and the negative binomial provided a significantly better fit to the data). To ensure all predictor variables were on the same scale, we applied a normal-quantile transformation to each variable prior to fitting the model. To select a model that best explains the data without including unnecessary variables, we used the “stepAIC” function to perform stepwise model selection via the Akaike information criterion (AIC). To assess correlations between predictor variables, we used ggplot2 v3.3.5 and the GGally v2.1.2 extension to create a scatterplot matrix and estimate Pearson correlation coefficients and p-values. To determine the impact of window-size on inferences about genomic correlates of discordance, we also generated genomic summary statistics (except D-statistics, which are unreliable for small windows; Martin et al. 2015) and performed multiple regression for 50 kb, 10 kb, and 5 kb windows. For the smaller window sizes (≤ 50 kb), we subsampled the data so that there was a 50 kb space between each window to avoid sampling tightly linked windows.

## Results

### Neodiprion De Novo *and Pseudo-Reference Genome Assemblies*

The *N. lecontei* iyNeoLeco1.1 assembly was sequenced to 93x coverage of PacBio HiFi reads and produced an assembly size of 272.074 MB in 106 scaffolds from 168 contigs with only 2% of the genome represented in gaps. Additional assembly statistics such as contig and scaffold N/L50 and N/L90 can be found in Table S2. Assessment for genome completeness using BUSCOs revealed that for all relevant databases from Eukaryota, Metazoa, Arthropoda, Insecta, Endopterygota, and Hymenoptera, genome completeness estimates ranged from 95.2% (Hymenoptera ortholog database v.10) to 99.6% (Eukaryota ortholog database v.10) (Table S3). All BUSCOs were found in assembled chromosomes, and none were in unplaced contigs.

For the remaining species, an average of 96.2% (range: 93.5-97.7%) of sequencing reads mapped to the *N. lecontei* reference genome, resulting in a final average coverage of 20.1x (range: 16x-28x) after removing regions with low or unusually high coverage (Table S4). The high mapping rates are likely attributable to low overall genetic divergence among eastern North American *Neodiprion* species: across all species, genome-wide average pairwise genetic distance (K80) was 0.0047 (range: 0.0003 - 0.0093) (Table S5).

### *Species-Tree Estimates for Eastern North American* Neodiprion

The *N. lecontei* genome and our 19 reference-based whole-genomes produced six windowed datasets, four gene datasets, and six SNP datasets, summarized in Table S6. Looking first at the species trees produced using a locus-based approach (ASTRAL-III), our “total evidence” (all data minus regions that did not pass our quality filters) analysis of windows produced strong support for 3 of 5 of species complexes: *lecontei, pinusrigidae*, and *pratti* (Fig. 2a). 5 of 6 members of the *virginianus* complex were recovered as monophyletic, but *N. dubiosus*—which was recovered as sister species to *N. rugifrons* in a previous analysis (Fig. 1)— clustered with 3 of 4 members of the *abbotii* complex. Identical topologies were recovered when the size of windows used to estimate gene trees was ≥ 10 kb, with declining quartet support as window size decreased. Only the 5 kb window analysis produced a different species tree, placing *N. compar* as sister to remaining *Lecontei* group species, with low quartet support throughout (Fig. 2b).

**Figure 2:**
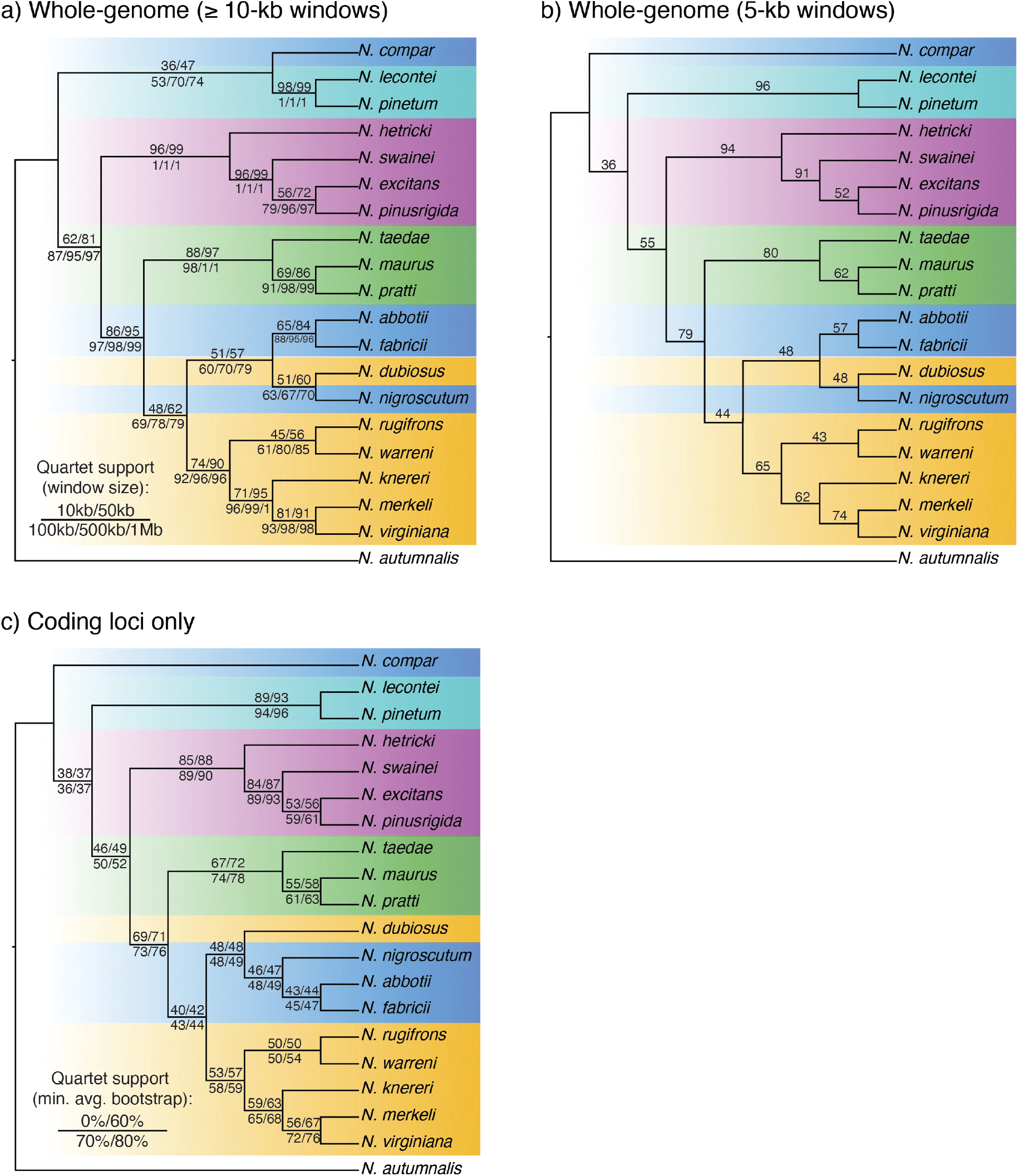
*Neodiprion* species trees estimated via ASTRAL-III. **a**) Consensus species-tree topology for whole-genome data divided into 10 kb, 50 kb, 100 kb, 500 kb, and 1000 kb windows, **b)** species tree for whole-genome data divided into 5 kb windows, and **c)** consensus species tree for analysis all coding regions with at least 10 parsimoniously informative sites, with no restrictions on gene-tree bootstraps as well as gene trees filtered to have a minimum average bootstrap value of 60%, 70%, or 80%. Colors correspond to the named species complexes in Fig. 1. Numbers on nodes indicate quartet support; for (**a**) and (**c**), support values are given in the order indicated in each figure legend.

As an alternative to estimating gene trees for adjacent windows, we isolated 11,969 aligned coding regions and produced a species tree using gene trees from 10,602 loci that had at least 10 parsimoniously informative sites (Fig. 2c). We also produced additional species trees using only gene trees that had a minimum average bootstrap value of 60%, 70%, or 80% (Fig. 2c; Table S6). All four coding-locus datasets produced identical topologies with minimal differences in quartet support. This topology differed from the large-window topology (Fig. 2a) only in the placement of *N. compar* and *N. dubiosus* and from the 5 kb window topology (Fig. 2b) only in the placement of *N. dubiosus*. Quartet support for the coding-locus tree was similar to that observed for the 5 kb window tree.

Turning to SNP-based (SVDquartets) species trees, our “total evidence” SNP dataset (all SNPs that passed quality and completeness thresholds) consisted of 13,732,314 SNPs (Table S6). This all-SNPs dataset and a dataset consisting of SNPs sampled every1 kb (212,496 SNPs) produced the same topology, both with high bootstrap support for all nodes except for a node that recovered *N. compar* as sister to remaining *Lecontei* group species (Fig. 3a). SNPs sampled at 5 kb and 10 kb intervals (46,473 SNPs and 12,427 SNPs, respectively) produced a topology that differed from the all-SNPs tree in the placement of the *pratti* group (Fig. 3b). Bootstrap support also tended to decline as the size of the SNP dataset decreased (Fig. 3). The topology became increasingly unstable for the smallest SNP datasets: 1 SNP per 50 kb (5,131 SNPs; Fig. 3c) and 1 SNP per 100 kb (2,611 SNPs; Fig. 3d).

**Figure 3:**
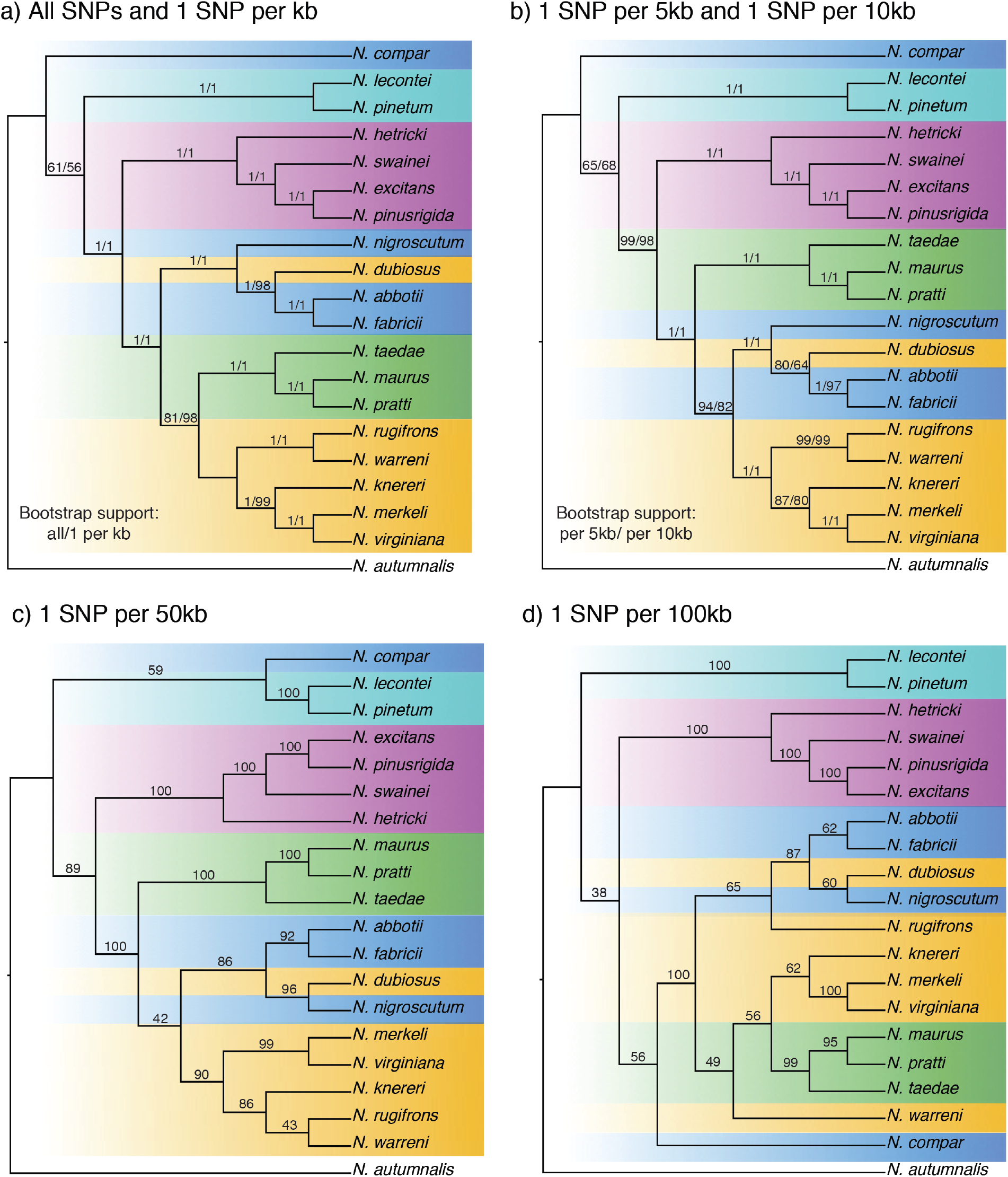
*Neodiprion* species trees estimated via SVDquartets. **a**) Consensus species tree for all SNPs and SNPs subsampled every 1,000 base pairs, **b**) consensus species tree for SNPs sampled every 5 kb or 10 kb, **c)** species tree for SNPs sampled every 50,000 bp, and **d)** species tree for SNPs sampled every 100 kb. Colors correspond to the named species complexes in Fig. 1. Values at each node are bootstrap support obtained using 10,000 bootstrap replicates in PAUP, with order in (**a**) and (**b**) as indicated in the figure legend.

Across all species trees (except the smallest SNP dataset), we consistently recovered five primary clades: *N. lecontei* + *N. pinetum* (*lecontei* species complex); *N. excitans* + *N. hetricki* + *N. pinusrigidae* + *N. swainei* (*pinusrigidae* species complex); *N. maurus* + *N. pratii* + *N. taedae* (*pratti* species complex), *N. abbotii* + *N. dubiosus* + *N. fabricii* + *N. nigroscutum* (*abbotii* species complex minus *N. compar* and plus *N. dubiosus*), and *N. knereri* + *N. merkeli* + *N. rugifrons* + *N. virginiana* + *N. warreni* (*virginianus* complex minus *N. dubiosus*) (Fig. 2 and 3). Most of the relationships within these clades were also highly consistent across analyses and datasets. However, three relationships were sensitive to sampling approach and analysis method: 1) the placement of *N. dubiosus* within the *abbotii* clade, 2) the placement of *N. compar* relative to remaining *Lecontei* group species, and 3) the placement of the *pratti* clade relative to the *virginianus* and *abbotii* clades.

### Genomic Patterns of Gene-Tree Discordance

To explore variation in gene-tree topology across the genome, we focused on the windowed datasets. First, we visualized the topologies estimated from 500 kb windows that had < 10% missing data using DensiTree (Fig. 4). Relationships that were insensitive to locus-sampling and analysis approach (Figs. 2 and 3) also had high levels of concordance across gene trees (Fig. 4). For example, the DensiTree plot revealed consistent topological support for the *lecontei, pinusrigidae*, and *pratti* complexes, as well as relationships within these complexes and within the *virginianus* complex (minus *N. dubiosus*). By contrast, there was considerable heterogeneity among gene trees in the placement of *N. compar*, relationships within the *abbotii* complex, and relationships between the *abbotii, pratti*, and *virginianus* complexes. Although heterogeneity is expected under incomplete lineage sorting, genome-wide D-statistics (bottom diagonal, Fig. 4) and mitochondrial gene flow estimates (2*Nm*) (top diagonal, Fig. 4) reveal evidence of introgression in areas of the tree with gene-tree discordance. For example, *N. rugifrons, N. dubiosus*, and *N. nigroscutum* have some of the highest genome-wide D-statistics and mitochondrial population migration rates (*N. dubiosus* vs. *N. rugifrons*: D = 0.32, mitochondrial 2*Nm* = 2.34; *N. nigroscutum* vs. *N. rugifrons*: D = 0.29, mitochondrial 2*Nm* = 12.55; *N. nigroscutum* vs. *N. dubiosus*: D = not estimated; mitochondrial 2*Nm* = 7.69). Notably, although the introgression metrics we evaluated used different data and analysis approaches, we nevertheless found that they two sets of estimates were significantly correlated (Spearman’s rho = 0.208, *P* = 0.0209).

**Figure 4:**
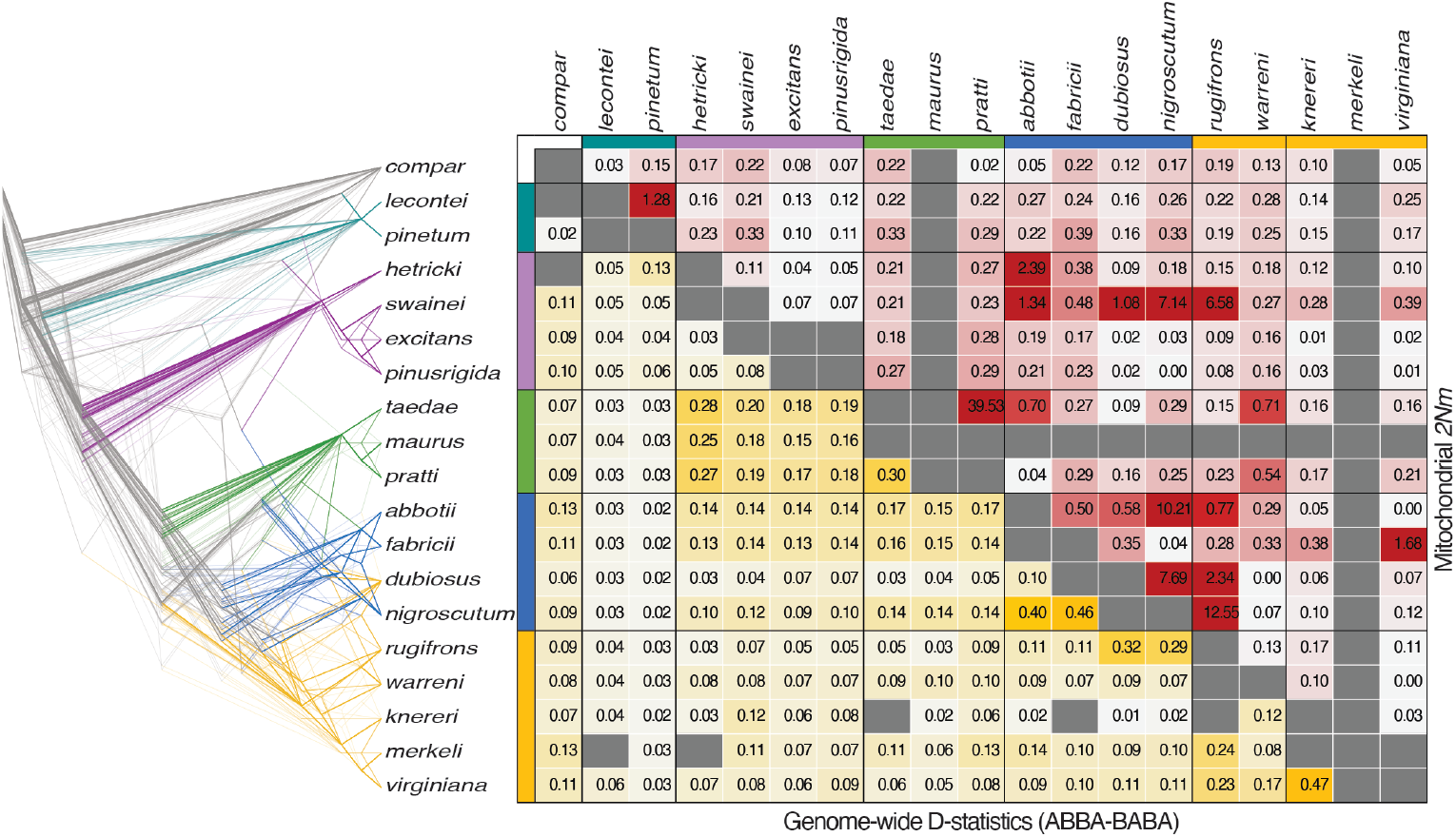
Genome-wide patterns of topological discordance and mitochondrial and nuclear introgression metrics. The tree on the left was generated using DensiTree and depicts topologies for 392 gene-trees estimated from 500 kb windows with < 10% missing data. The lines and colored bars next to species names are colored according to named species complexes presented in Fig. 1. The lower diagonal (yellow) represents genome-wide average D-statistics for each species pair, with deeper yellow signifying more evidence for nuclear introgression. Note that because one exemplar is sampled per species, D-statistics cannot be calculated for sister species, and these missing values are indicated in gray. The upper diagonal (red) represents estimates of mitochondrial gene flow (2*Nm*) that were calculated using coalescent-based methods and multiple individuals per species (Linnen and Farrell 2007) with redder colors signifying higher estimated rates of mitochondrial introgression. Note that 2*Nm* values could not be calculated for comparisons involving *N. maurus* and *N. merkeli* because only a single individual was available for each species (Linnen and Farrell 2007); these missing values are indicated in gray.

To quantify the overall extent of gene-tree discordance, we calculated the Robinson-Foulds distance between each window-based gene tree from each dataset (1 Mb, 500 kb, 100 kb, 50 kb, 10 kb, and 5 kb) and the window-based species tree (Fig. 2a). Regardless of dataset, most gene trees did not match the inferred species tree (Table S7). Overall, the proportion of gene trees matching the species trees increased as the window size increased, from 0.00007 in the 5 kb window dataset to 0.33 in the 1 Mb window dataset. Similarly, the maximum observed Robinson-Foulds distance was highest in the 5 kb window datatset (RF = 34) and lowest in the 1 Mb dataset (RF = 18).

To visualize how gene-tree discordance is distributed across the genome, we painted the chromosomes according to the Robinson-Foulds distance between the gene tree for each locus and the reference species tree (Fig. 5). Regardless of the window size used to paint the chromosomes, we observed both high-concordance (low Robinson-Foulds distance) and low-concordance (high Robinson-Foulds distance) regions of the genome. As window size decreased, variation in topology increased overall. Regardless of window size, we found a cluster of unusually high-discordance gene-trees located on Chromosome 1 between 27 Mb and 30 Mb (Fig. 5).

**Figure 5:**
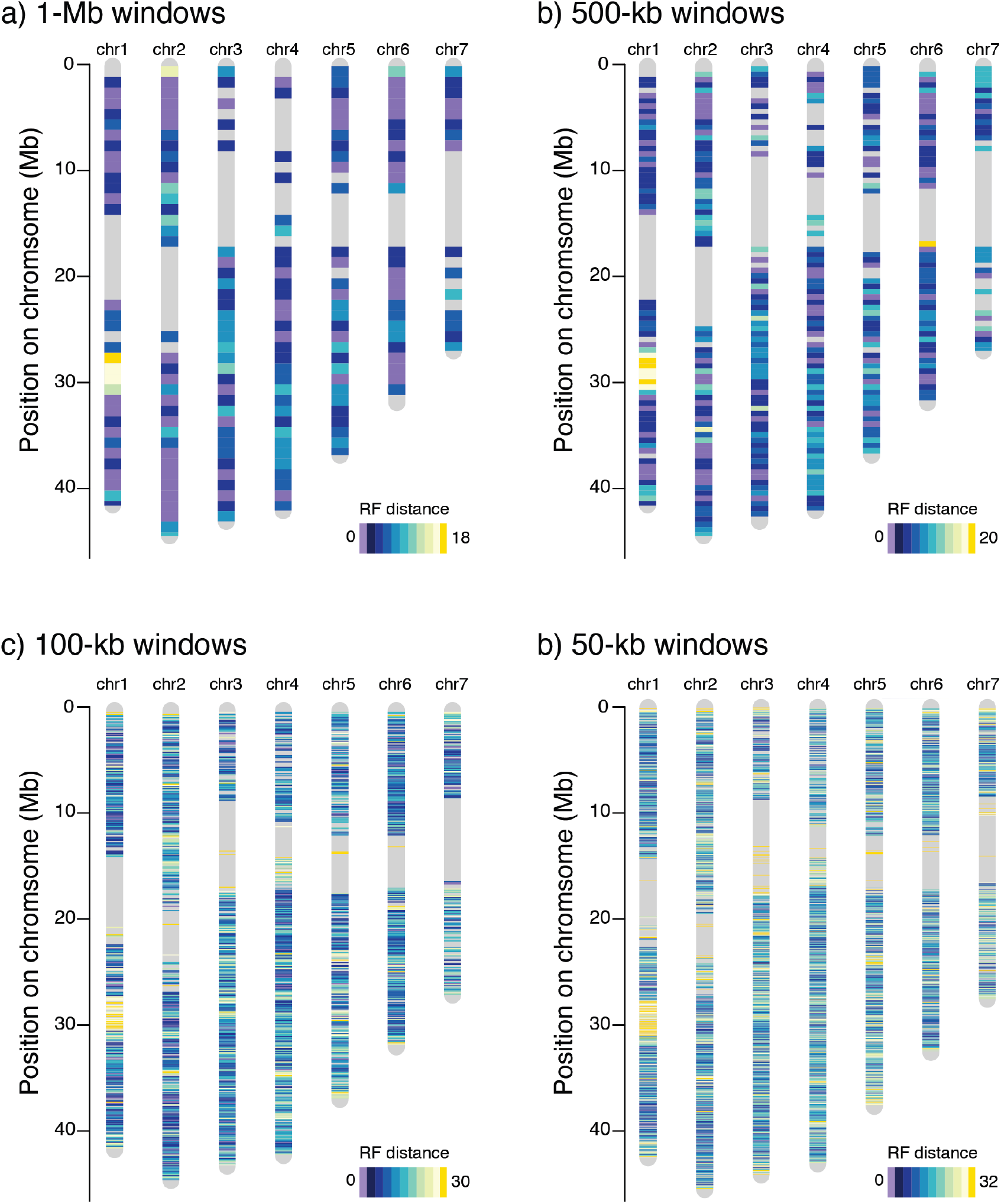
Genomic landscape of gene-tree discordance for different window sizes. Each plot depicts the seven chromosomes in the *Neodiprion* genome, with position in Mb on the y-axis. Each band represents the Robinson-Foulds (RF) distance between the gene tree for that window and the species tree estimated from the windowed datasets (Figure 2A). Discordance patterns are shown for **a)** 1000 kb windows, **b)** 500 kb windows, **c)** 100 kb windows and **d)** 50 kb windows. The colors of the bands are a purple to gold gradient with purple bands representing an exact match to the species tree (RF distance = 0), dark blue representing trees that are very similar to the species tree, and gold representing the most divergent trees. Regions that were excluded based on missing data (> 10%)—mostly in centromeric and telomeric regions—are colored in gray.

To explore factors that give rise to heterogeneous discordance across the genome, we estimated genomic variables for each of the 500 kb, 50 kb, 10 kb, and 5 kb windows for which we calculated Robinson-Foulds distances from the species trees (Tables S8-S11). To estimate local recombination rate, we constructed a new genetic map using sequencing reads from a previous cross between *N. lecontei* populations (Linnen et al. 2018). By mapping reads to a new reference genome, we nearly tripled the number of markers (from 503 to 1436) and decreased the average spacing between markers from 2.4 cM (maximum spacing = 24.3 cM) to 0.9 cM (maximum spacing = 13.2 cM). The total map length was 1271.6 cM. With an assembled genome size of ∼272 Mb, this corresponds to a genome-wide recombination rate of 4.68 cM/Mb, a slightly higher rate than was previously reported (3.43 cM/Mb; Linnen et al. 2018). Plotting genetic distance as a function of physical distance revealed an even coverage of markers across chromosomes; gaps in the new genetic map corresponded to low-recombination centromeric regions (Figs. S2-S3). Using these data, local recombination rates were estimated via sliding windows, revealing heterogeneous recombination rates across the genome (Figs. S2-S3).

We also investigated pairwise correlations between all genomic predictor variables (Fig. S4) and found that the sign of the correlations was generally robust to window sizes, although smaller windows (more data points) recovered more significant correlations (Table S12). Overall, the average bootstrap value of gene trees correlated positively with the number of parsimony informative sites, gene density, and average genetic distance; and negatively with GC content. Average genetic distance also correlated positively with number of parsimony informative sites and gene density, and negatively with recombination rate and GC content. Recombination rate and GC content were positively correlated across all datasets (Table S12).

Across the four datasets (500 kb, 50 kb, 10 kb, and 5 kb windows), discordance (Robinson-Foulds distance) increased with gene density and average genetic distance, while discordance was negatively correlated with recombination rate and average bootstrap support (Table 1; Fig. 6). Discordance decreased as the number of parsimony informative sites increased for all datasets except the 500 kb windows (Table 1). GC content was positively correlated with discordance for the 500 kb dataset (Fig. 6b), but negatively correlated with discordance for the 10 kb dataset (Table 1). However, GC content for the 500 kb dataset was approximately double that of the smaller-window datasets (Table S7), likely due to the exclusion of GC-poor regions at the telomeres and centromeres for the large-window dataset (Fig. 4). We only estimated D-statistics for the largest windows (500 kb) and found that higher average D-statistics were associated with higher discordance, although this relationship was weaker than for other genomic variables (Table 1; Fig. 6).

**Table 1.** Results of negative binomial regression and stepwise model selection for predicting Robinson-Foulds distances between gene trees and whole-genome species tree for four different window sizes (500 kb, 50 kb, 10 kb, and 5 kb).

| Window Size | Variable | Coefficient | Std. Error | z-value | Pr(>\|z\|) |
| --- | --- | --- | --- | --- | --- |
| 500 kb | (Intercept) | 1.221 | 0.0480 | 25.45 | < 2e-16 |
|  | Average bootstrap support | -0.141 | 0.0483 | -2.91 | 0.003577 |
|  | GC content | 0.154 | 0.0594 | 2.59 | 0.009512 |
|  | Average pairwise genetic distance (K80) | 0.334 | 0.0629 | 5.30 | 1.14E-07 |
|  | Average D-statistic | 0.076 | 0.0481 | 1.58 | 0.114077 |
|  | Gene density | 0.109 | 0.0520 | 2.09 | 0.036398 |
|  | Recombination rate (cM/MB) | -0.163 | 0.0483 | -3.37 | 0.000756 |
| 50 kb | (Intercept) | 2.074 | 0.0122 | 170.32 | < 2e-16 |
|  | # parsimony-informative sites | -0.187 | 0.0175 | -10.69 | < 2e-16 |
|  | Average bootstrap support | -0.037 | 0.0126 | -2.97 | 0.00302 |
|  | Average pairwise genetic distance (K80) | 0.212 | 0.0179 | 11.87 | < 2e-16 |
|  | Gene density | 0.081 | 0.0137 | 5.93 | 3.08E-09 |
|  | Recombination rate (cM/MB) | -0.097 | 0.0122 | -7.95 | 1.81E-15 |
| 10 kb | (Intercept) | 2.608 | 0.0053 | 492.91 | < 2e-16 |
|  | # parsimony-informative sites | -0.097 | 0.0078 | -12.49 | < 2e-16 |
|  | Average bootstrap support | -0.019 | 0.0055 | -3.49 | 0.000489 |
|  | GC content | -0.021 | 0.0060 | -3.42 | 0.000629 |
|  | Average pairwise genetic distance (K80) | 0.090 | 0.0079 | 11.41 | < 2e-16 |
|  | Gene density | 0.026 | 0.0057 | 4.56 | 5.23E-06 |
|  | Recombination rate (cM/MB) | -0.056 | 0.0054 | -10.48 | < 2e-16 |
| 5 kb | (Intercept) | 2.808 | 0.0043 | 650.68 | < 2e-16 |
|  | # parsimony-informative sites | -0.041 | 0.0066 | -6.15 | 7.55E-10 |
|  | Average bootstrap support | -0.041 | 0.0048 | -8.55 | < 2e-16 |
|  | Average pairwise genetic distance (K80) | 0.035 | 0.0062 | 5.63 | 1.82E-08 |
|  | Gene density | 0.012 | 0.0045 | 2.61 | 0.00904 |
|  | Recombination rate (cM/MB) | -0.043 | 0.0044 | -9.80 | < 2e-16 |

**Figure 6:**
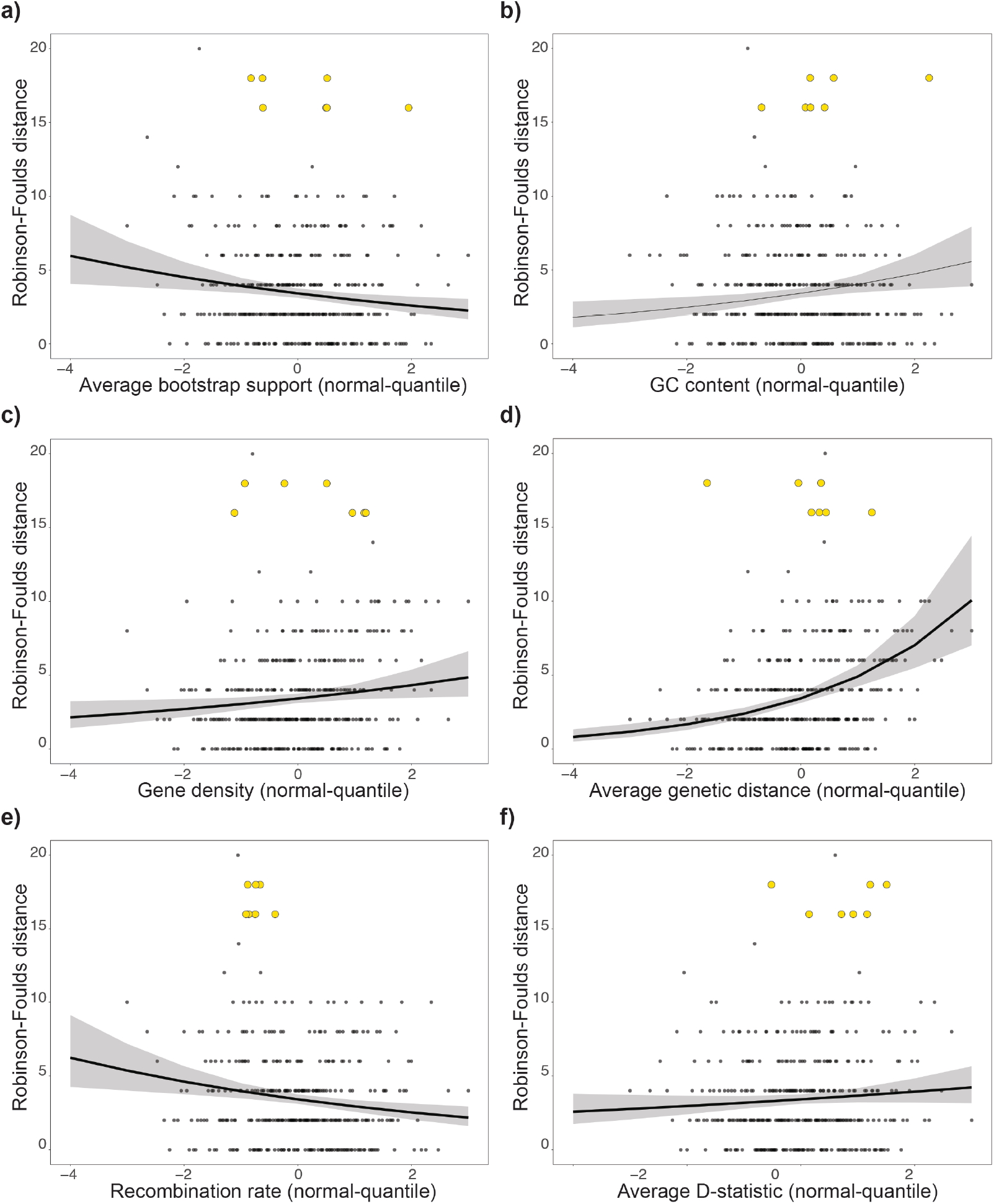
Genomic correlates of gene-tree discordance for 500 kb windows. Each panel depicts the Robinson-Foulds (RF) distance between each 500 kb gene tree and the species tree in Fig. 2a as a function of one of six variables that were retained after stepwise model selection (Table 1), including: **a)** average bootstrap support across all nodes in each gene tree, **b)** GC content, **c)** gene density, **d)** average pairwise genetic distance (K80), **e)** local recombination rate estimated via sliding window (cM/Mb), and **f)** average D-statistic. All predictor variables are normal-quantile transformed. Curves on each plot give the predicted values for RF distance as a function of genomic variable values (marginal effects) estimated via negative binomial regression and accounting for other variables in the model. The seven yellow points on each plot correspond to the seven 500 kb windows located in the discordance hotspot on Chromosome 1 (Fig. 5).

Although several genomic variables predicted gene-tree discordance, these parameters do not appear to explain the cluster of adjacent windows with unusually high discordance on Chromosome 1 (gold bars in Fig. 5; yellow dots in Fig. 6). Windows in this region appear to have slightly higher D-statistics and lower recombination rates than the genome-wide average (all quantile-normalized values for yellow dots > 0 and < 0 in Figs. 6f and 6e, respectively). However, Robinson-Foulds distances in the Chromosome 1 discordance “hotspot” are much higher than windows with comparable values for all genomic predictors (Fig. 6). The window with the highest Robinson-Foulds distance from the species tree (RF = 20) is located within the centromeric region of Chromosome 6 (Fig. 5b) and is on the low-end for estimates average bootstrap support (Fig. 6a).

Because genomic predictors do not appear to explain the discordance hotspot on Chromosome 1, we investigated this region further. First, we mapped diapause strategy and preferred pine host—two traits that are thought to contribute to reproductive isolation (Knerer and Atwood 1973; Linnen and Farrell 2010; Bendall et al. 2017)—onto the consensus species tree derived from all “total evidence” (all windows, all SNPs) analyses (Fig. 7a). Then, we constructed a consensus tree for the seven gene trees estimated from the discordance hotspot to see which species or clades had unusual placements (Fig. 7b). Compared to the whole-genome species tree (Fig. 7a), the consensus gene tree for the discordance hotspot (Fig. 7b) recovered closer relationships among egg-overwintering species (in *pinusrigidae* and *pratti* groups) and among jack-pine feeders in the *abbotii* and *virginianus* clades (*N. abbotii, N. nigroscutum, N. dubiosus*, and *N. rugifrons*). The discordance hotspot tree also has some striking similarities to the mitochondrial gene tree estimated in Linnen and Farrell 2007 (Fig. 7c): both trees place the *pratti* complex plus *N. warreni* with members of the *pinusrigidae* complex and cluster members of the *abbotii* (minus *N. compar*) and *virginianus (*minus *N. warreni*) complexes into a single clade. Going a step further, we used custom scripts and the NCBI *N. lecontei* annotation to determine which genes were present in the discordance hotspot. In total, we identified 203 genes in this 3.5-Mb region, including eight genes potentially involved in chemosensation (putative gustatory, olfactory, and ionotropic receptors), nine genes potentially involved in mito-nuclear interactions, and six genes potentially related to pre- and post-copulatory traits (Table S13).

**Figure 7:**
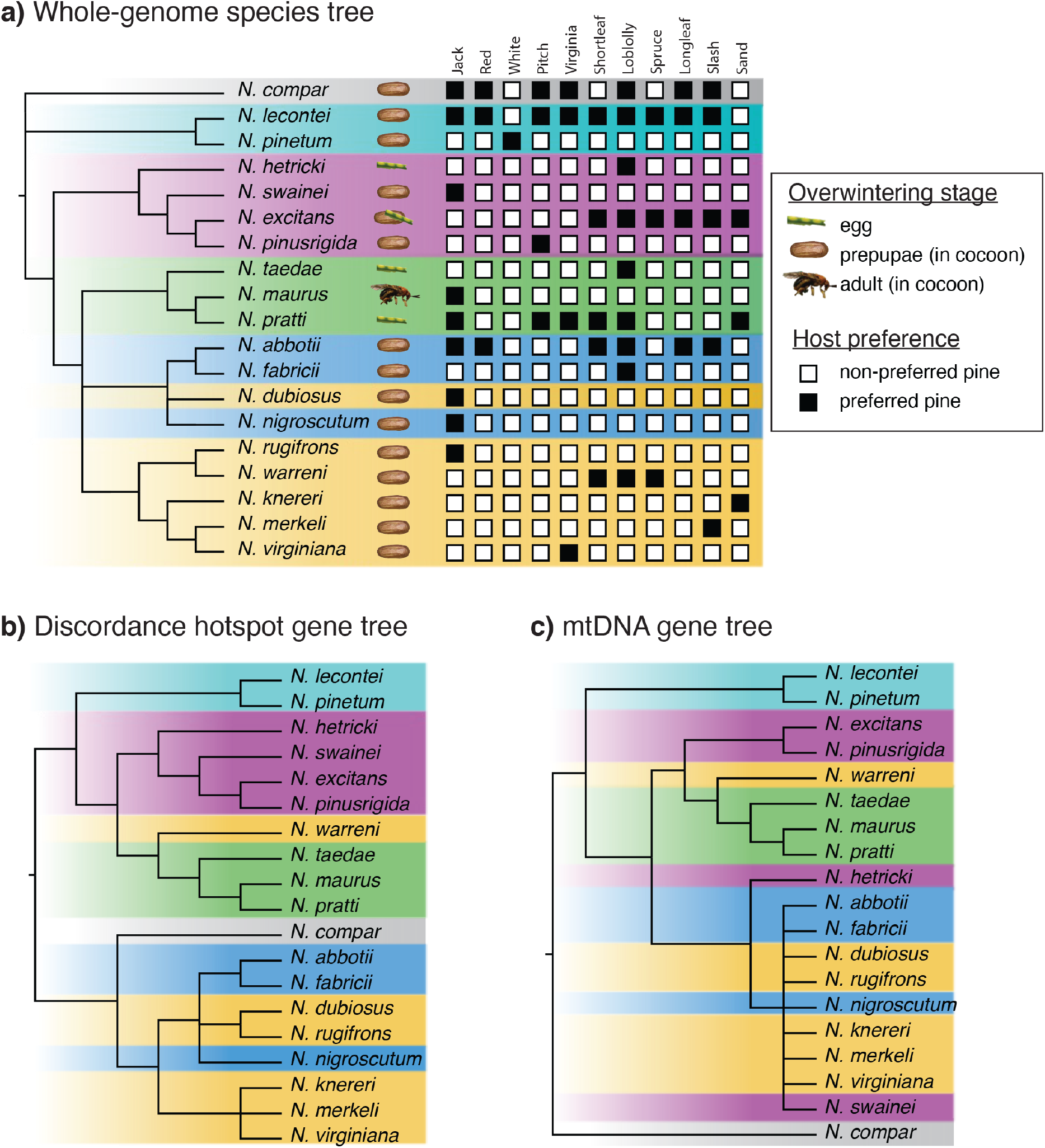
Exploration of mechanisms generating the Chromosome 1 discordance hotspot. **a)** consensus species tree produced by whole-genome data and two species-tree methods (Figs 2 and 3), **b)** consensus gene tree for the 3.5 Mb discordance hotspot on Chromosome 1, and **c)** collapsed mitochondrial gene tree (modified from Linnen and Farrell 2007, collapsed such that each species is represented by a single tip). In **(a)**, three distinct overwintering strategies are represented by images. Host use is indicated by a matrix of black and white boxes, with black indicating that a pine is a preferred host and white indicating a pine is not a preferred host (modified from Linnen and Farrell 2010).

## Discussion

Whole-genome datasets provide a comprehensive sample of heterogeneous histories, and the multi-species coalescent provides a framework for modeling sources of heterogeneity. Here, we combine a high-quality reference genome with whole-genome alignments for 20 pine sawfly species to achieve three goals: 1) determine the effect of sampling strategy and analysis method on species-tree estimation from whole-genome data, 2) estimate a species tree for eastern North American *Neodiprion* species, and 3) identify genomic correlates of gene-tree discordance. These analyses also revealed an approximately 3.5-Mb region on Chromosome 1 that generates gene trees with especially high discordance from the estimated species tree. Below, we discuss progress on each of our three goals and possible explanations for the discordance hotspot, while highlighting both the broader implications and limitations of our analyses.

### Impact of Locus-Sampling and Analysis Method on Species-Tree Reconstruction

Although there is growing body of research investigating how marker sampling strategies and species-tree analysis methods impact accuracy of species-tree estimates, many outstanding questions remain (Chou et al. 2015; Chen et al. 2017; Reddy et al. 2017; Huang et al. 2020; Karin et al. 2020; Alda et al. 2021; Literman and Schwartz 2021; Mongiardino Koch 2021). In the absence of a consensus for phylogenomic best practices, whole-genome alignments offer an opportunity to investigate how partitioning and analyzing the data in different ways affect species-tree estimates. Our analysis of whole-genome alignments from closely related pine sawfly species revealed several general patterns that have broader implications for phylogenomic analysis of recently diverged lineages.

First, we found that ASTRAL analyses of gene trees generated from chromosomal windows ranging from 10 kb to 1000 kb produced identical species-tree topologies (Fig. 2a). This observation is consistent with simulation studies that demonstrate that intralocus recombination has a minimal impact on the accuracy of species-tree inference (Lanier and Knowles 2012; Wang and Liu 2016; Zhu et al. 2022). Given that a 100-fold difference in window size had no impact on the inferred species-tree topology, our finding that a two-fold difference in window size (5 kb vs. 10 kb) resulted in slightly different species-tree topologies (Fig. 2b) is unlikely to be explained by differences in opportunities for intralocus recombination. Instead, since shorter loci tend to have less phylogenetic signal, differences between species trees estimated from 5 kb and 10 kb gene trees may reflect increased gene-tree estimation errors for the shorter 5 kb loci (Betancur-R. et al. 2014; Blom et al. 2016; Huang et al. 2020; Mongiardino Koch 2021; but see Chou et al. 2015). Beyond topology, another difference observed among species trees produced by different window sizes was that quartet support tended to decline as window size decreased (Fig. 2). This is unsurprising since larger windows are more likely to sample multiple genealogical histories (Degnan and Rosenberg 2009), thereby masking gene-tree heterogeneity that could decrease quartet support. However, there were some nodes that had relatively low quartet support (< 80) regardless of window size. These low-support nodes tended to be those that were most sensitive to species-tree method (e.g., clades containing *N. dubiosus* and *N. compar*), indicating that even though constructing gene trees from large windows may mask local genealogical heterogeneity, this heterogeneity is still being captured genome-wide and reflected in nodal support.

Second, we found that subsampling coding sequences from the whole-genome dataset—a dataset analogous to one that might be produced from transcriptome sequencing or exome capture—had only a modest impact on the inferred species tree (Fig. 2c vs. Fig. 2a and Fig. 2b). Compared to the whole-genome windowed dataset, a dataset of 10,601 genes (coding sequences only) averaging 3,335 base pairs in length differed from the 10-1000 kb window species tree in the placement of two species (*N. compar* and *N. dubiosus*) and from the 5 kb window species tree in the placement of one species (*N. dubiosus*). Although our protein-coding genes are contained within the genomic windows, most of the windowed data consists of intergenic or intronic sequence. Thus, our finding that chromosomal windows (largely non-coding) and genes (coding sequence only) produced slightly different topologies is consistent with previous work that indicates that data type (coding vs. non-coding sequence) can influence species-tree estimates (Chen et al. 2017; Reddy et al. 2017; Alda et al. 2021; Literman and Schwartz 2021). Possible reasons why coding regions are more likely to produce discordant gene trees than non-coding regions—thereby producing conflicting species trees—are considered below (see “*Genomic Predictors of Gene-Tree Discordance*”). That said, topological conflicts between the windowed dataset and the protein-coding dataset for this recently diverged clade (< 10 mya) were modest in comparison to more striking differences that have been documented in more distantly related taxa (e.g., hundreds of millions of years; Reddy et al. 2017; Literman and Schwartz 2021). Thus, the impact of data type on species-tree inference may be dependent on the divergence times of the taxa under study.

Third, we found that SNP-based species-tree analysis is very sensitive to how much of the genome is sampled. Like the locus-based ASTRAL analyses, the placement of *N. compar* and *N. dubiosus* in SVDquartets trees varied depending on how the data were sampled (Fig. 3). In addition, the placement of the *pratti* complex and relationships among species in the *virginianus* complex were affected by the size of the SNP dataset. Except for low bootstrap support for the placement of *N. compar*, the largest SNP datasets had high bootstrap support for all branches (Fig. 3a). This contrasts with the locus-based approach that had reduced nodal support for multiple branches that were sensitive to alternative sampling strategies (Fig. 2). That relationships are unstable with reduced SNP number is consistent with simulation work demonstrating that under some parameter combinations, large numbers of SNPs may be needed for accurate species-tree inference using this method (Long and Kubatko 2018; Wascher and Kubatko 2021). In our case, it was necessary to sample hundreds of thousands to millions of SNPs (Table S6) to obtain a stable, highly supported species-tree estimate using SVDquartets. Although SVDquartets assumes that SNPs are independent—an assumption that is almost certainly violated in our all-SNPS and 1 kb-SNPs datasets—previous work demonstrates that this method performs well even for multilocus data with linked sites (Chifman and Kubatko 2014; Chou et al. 2015). Notably, our smaller SNP datasets (∼2,000-15,000 SNPs) were on par with those that might be generated via RADseq coupled with choosing a single site per RAD locus. Based on our results and previous studies of the behavior of SVDquartets, phylogenomic studies using RADseq should choose library preparation protocols and filtering strategies that maximize the number of loci and SNPs available for analysis.

A final pattern revealed by our analyses was that locus-based (Fig. 2) and SNP-based (Fig. 3) approaches produced identical results for most branches in the species tree, except for the same three lineages (*N. compar, N. dubiosus*, and the *pratti* species complex) that were sensitive to locus/SNP sampling within each approach. For *N. dubiosus* and the *pratti* species complex, high D-statistics and mitochondrial introgression estimates indicate that at least some of the topological uncertainty for these lineages stems from post-divergence gene flow (Fig. 4). More generally, interspecific gene flow has been pervasive in the *Lecontei* group (Linnen and Farrell 2007; Bendall et al. 2022; Fig. 4), a scenario for which SVDquartets may outperform ASTRAL and other locus-based approaches (Long and Kubatko 2018; Wascher and Kubatko 2021). Conversely, there is less evidence of gene flow between *N. compar* and other *Lecontei* group species (Fig. 4), suggesting that topological uncertainty for this species may stem from incomplete lineage sorting, a scenario for which ASTRAL may outperform SVDquartets (Chou et al. 2015). Thus, one explanation for lineages that were not placed consistently is that they diverged under scenarios that are particularly challenging for ASTRAL or SVDquartets, including pervasive incomplete lineage sorting, introgression, or both. Alternatively, unstable relationships across methods could result from simultaneous divergence of more than two descendant lineages from a single ancestor, a scenario commonly referred to as a “hard polytomy” (Maddison 1989).

While we have explored two types of species-tree methods (locus-based and SNP-based) and several different ways of partitioning whole-genome alignments, we have not exhaustively compared all species-tree methods or locus-sampling strategies. For example, data could be further subsampled to mimic other types of phylogenomic datasets that use different types of markers (e.g., UCEs). Although our whole-genome alignment would be prohibitive for full-likelihood methods, subsampled data could be analyzed using full-likelihood methods that account for uncertainty in gene tree estimation, as well as sources of discordance other than incomplete lineage sorting. A key question, however, is whether the size of datasets that could be analyzed in a reasonable time frame would be sufficient for the complex models under consideration. Consideration of divergence models that include hybrid species formation—which has been hypothesized for at least one *Neodiprion* species (Ross 1961)—may also be informative (e.g., Blischak et al. 2018). Finally, we have analyzed only a single exemplar per species, and previous work on the *Lecontei* clade (albeit with far fewer loci) suggests that taxon sampling can have a large effect on the inferred species tree (Linnen and Farrell 2008a). Also, when multiple individuals are sampled per species, a suite of complementary methods can be used to estimate demographic parameters, introgression rates, and population structure that might impact species-tree inference. For these reasons, examining the impact of taxon sampling is also a high priority for future work on the *Lecontei* species group.

### An Updated Lecontei Group Species Tree

Aside from three branches with uncertain placement, most relationships among *Lecontei* group species were recovered with high support across methods and datasets (Fig. 2, 3). Our updated species tree (Fig. 7a) resembled the previous three-locus species tree (Fig. 1) but recovered an additional Ross (1955) species complex (*pratti* group) and resolved most relationships within species complexes with high support. Both old and new species trees strongly reject the placement of *N. compar* within the *abbotii* complex, a relationship that was proposed by Ross (1955) based on similar larval coloration and behavior (Fig. 1). Thus, our data suggest that similarities between *N. compar* and remaining members of the *abbotii* complex— which tend to be more solitary and cryptically colored than other species—are due to phenotypic convergence. Based on these results, we propose an updated *N. abbotii* complex that excludes *N. compar*.

Despite some uncertainty in the placement of *N. dubiosus*, our analyses consistently placed this species within the *abbotii* species group, rendering the *virginianus* complex polyphyletic and the *abbotii* complex paraphyletic. Curiously, the previous three-gene species tree—which contained multiple *N. dubiosus* individuals—consistently placed *N. dubiosus* within the *virginianus* species complex and recovered the *abbotii* complex (minus *N. compar*) as monophyletic (Linnen and Farrell 2007, 2008a). This discrepancy could simply mean that the whole-genome data enabled us to recover a more accurate species-tree estimate. However, *N. dubiosus* adults are nearly indistinguishable from *N. rugifrons* adults (Becker et al. 1966), raising the possibility that placement within the *abbotii* complex in the whole-genome tree is incorrect. For example, it is possible that the *N. dubiosus* exemplar we chose, which had typical *N. dubiosus* morphology, had an admixed genome due to recent hybridization. We therefore refrain from reassigning *N. dubiosus* to the *abbotii* complex until additional samples can be analyzed.

### Genomic Predictors of Gene-Tree Discordance

Although most nodes in our species-tree topology were robust to sampling and analysis method, there was nevertheless substantial gene-tree heterogeneity across the genome (Fig. 5,6), a pattern observed in many other taxa that diverged rapidly, often with gene flow (Pease et al. 2016; Edelman et al. 2019; Alda et al. 2021; Kozak et al. 2021; Meleshko et al. 2021; Zhang et al. 2021). In fact, regardless of the window size used, most of the inferred gene trees did not match the ASTRAL species tree (Table S7). Our high-quality chromosome-level assembly provided us with an opportunity to investigate mechanisms that give rise to heterogeneous levels of discordance across the genome. Because several genomic predictors of topological discordance are correlated (Fig. S4; Table S12), we used a multiple regression approach to disentangle the contributions of individual variables to discordance while controlling for other variables. We found that all seven genomic variables predicted topological discordance between gene trees and a reference species tree, although the effects of some variables were dependent on window size (Table 1, Fig. 6).

First, we found that the number of parsimony informative sites was negatively correlated with discordance (Robinson-Foulds distance from the species tree) for all but the 500 kb windows dataset (Table 1). This is perhaps unsurprising given that 500 kb windows contain the highest number of informative sites (Table S7). Second, for all window sizes, we found that gene trees with lower average bootstrap support tended to be more discordant from the estimated species tree (i.e., higher Robinson-Foulds distances; Fig. 6a, Table 1). This observation suggests that gene-tree estimation error contributes to genomic heterogeneity in gene-tree discordance. Three potential sources of gene-tree estimation error are: (1) a lack of phylogenetic signal (Rasmussen and Kellis 2007; Betancur-R. et al. 2014), (2) poor fit to nucleotide substitution models used to estimate gene trees (Reddy et al. 2017; Karin et al. 2020), and (3) mixed genealogical histories (Posada and Crandall 2002; Degnan and Rosenberg 2009). The first explanation—a lack of phylogenetic signal—likely applies only to the smaller windows. This argument is supported by a strong correlation between the number of parsimony informative sites and average bootstrap support for the 5 kb and 10 kb window datasets, but not the 50 kb and 500 kb window datasets (Table S12).

By contrast, misspecification of nucleotide substitution models is likely to be an issue for all window sizes. Although we allowed IQ-tree to choose the best-fit model for each locus, we did not partition loci to allow for different substitution models within loci, which is likely to be especially problematic when gene trees are estimated from large genomic windows. Moreover, standard nucleotide substitution models may be inadequate for describing complex patterns of substitution, particularly in coding regions (Whelan and Goldman 2001; Yang and Nielsen 2002; Whelan 2008; Chi and Liberles 2016; Echave et al. 2016; Reddy et al. 2017; Dornburg et al. 2019; Karin et al. 2020). Therefore, if model misspecification contributes to gene-tree discordance, discordance would be expected to be highest in gene-rich regions of the genome, which is exactly what we observed in our data: gene density is positively correlated with Robinson-Foulds distance for all window sizes (Table 1, Fig. 6c).

One widespread phenomenon that may not be sufficiently modeled in most standard nucleotide substitution models is heterogeneity in base composition among loci and taxa (Phillips et al. 2004; Romiguier and Roux 2017). An important driver of base compositional heterogeneity is GC-biased gene conversion, which is most pronounced in high-recombination regions of the genome (Eyre-Walker 1993; Galtier et al. 2001; Duret and Galtier 2009; Pessia et al. 2012; Figuet et al. 2015). Further complicating gene-tree inference in GC-rich regions, repeated bouts of GC-biased gene conversion and compensatory mutations are expected to lead to high levels of homoplasy (Jarvis et al. 2014; Romiguier and Roux 2017). Consistent with patterns in numerous taxa (e.g., Fullerton et al. 2001; Backström et al. 2010; Stevison and Noor 2010; Roesti et al. 2013), we found a positive correlation between GC content and recombination rate for all window sizes (Fig. S4, Table S12). Moreover, like many other phylogenomic datasets (Jarvis et al. 2014; Bossert et al. 2017; Reddy et al. 2017; Romiguier and Roux 2017), we found that GC content was positively correlated with gene-tree discordance, but only for our largest window size (Table 1, Fig. 6b). GC content was not retained in the stepwise regression models for either the 5 kb or the 50 kb datasets. For the 10 kb window dataset, GC content was *negatively* correlated with gene-tree discordance (Table 1). While this may seem surprising, we suspect the reason that the correlation between GC content and discordance was negative or non-existent when using smaller windows is that more loci in GC-poor telomeres and centromeres passed our missing-data filter for the smaller windows. These regions are likely to be highly repetitive and lack phylogenetic signal, a pattern that can be seen in our chromosome plots: as window size decreases, more regions are recovered at the ends and the middle of the chromosome, and these almost always produce highly discordant gene trees (Fig. 5). Furthermore, average GC content is much lower for the 5 kb, 10 kb, and 50 kb windows than for the 500 kb windows (Table S7).

In addition to a lack of phylogenetic signal and model misspecification, a third source of gene-tree estimation error—and therefore discordance—is the number of distinct genealogical histories that are sampled in a locus of a fixed size, which should correlate positively with recombination rate. While intralocus recombination appeared to have essentially no effect on the estimated species tree (Fig. 2a), it may nevertheless affect patterns of gene-tree discordance across the genome. For example, using simulated recombinant sequence alignments, Posada and Crandall (2002) demonstrated that when there is recent recombination between distantly related taxa (e.g., following hybridization and introgression), phylogenetic analysis can produce gene-tree estimates that differ from any of the histories within the alignment. Under this scenario, a positive association between recombination rate and gene-tree discordance might be expected since loci with high recombination rates will sample more histories. Alternatively, ancient recombination or recombination between closely related taxa tend to produce topologies corresponding to whatever genealogical history describes most of the positions in the locus (Posada and Crandall 2002). In this case, sampling more histories within a locus—especially a large locus—may converge on the most common evolutionary history and either have no effect on gene-tree discordance or even reduce discordance via masking aberrant, low-frequency gene-tree topologies. The negative relationship between gene-tree discordance and recombination rate observed in our data is consistent with this second scenario (Fig. 6e, Table 1). That said, under conditions expected to exacerbate deep coalescence, large loci with high intralocus recombination rates—the equivalent of concatenating many independent loci—may be particularly prone to converging on a gene-tree topology that does not match the species tree (Roch and Steel 2015).

A genome-wide association between recombination rate and discordance could also arise via pervasive linked selection. Because recombination reduces linked selection and increases local *N*_*e*_ (Smith and Haigh 1974; Charlesworth et al. 1993), discordance via incomplete lineage sorting is expected to correlate positively with recombination rate (Pease and Hahn 2013). While there is some empirical support for this prediction in primates and *Drosophila* (Hobolth et al. 2011; Prüfer et al. 2012; Pease and Hahn 2013), our regression models revealed that gene-tree discordance tended to be highest in ***low*** recombination regions of the genome after controlling for other genomic variables (Fig. 6e; Table 1). As discussed above, one potential explanation for reduced discordance in high-recombination regions is a local averaging effect that reduces the impact of low-frequency, highly discordant gene trees. A non-mutually exclusive explanation for high discordance in low-recombination regions is that positive selection increases gene-tree estimation error via both nucleotide model misspecification and homoplasy (Edwards 2009; Castoe et la. 2009). Due to linked selection (hitchhiking), the effects of positive selection—and any resulting gene-tree estimation error that contributes to discordance—will be most pronounced in low-recombination regions of the genome. Moreover, when divergent selection and is accompanied by gene flow, divergently selected alleles will tend to occur in low-recombination regions of the genome (Bürger and Akerman 2011; Yeaman and Whitlock 2011; Aeschbacher et al. 2017; Samuk et al. 2017). Thus, divergence-with-gene-flow would be expected to exacerbate associations between reduced recombination and increased discordance due to gene-tree estimation error.

Additional genomic patterns also support the hypothesis that positive selection and hitchhiking contribute to gene-tree estimation error and discordance in *Neodiprion*. For example, recombination rate is negatively correlated with average pairwise genetic distance, regardless of window size (Table S12). A genome-wide negative correlation between recombination rate and genetic divergence—a pattern unique to scenarios that include both divergent selection and gene flow (Nachman and Payseur 2012; Aeschbacher et al. 2017)—has been reported in several taxa (Keinan and Reich 2010; Geraldes et al. 2011; Brandvain et al. 2014; Samuk et al. 2017). Here, we show that this pattern is detectable across a clade of 19 species, despite using recombination rates that were estimated from a single species (*N. lecontei*) and clade-wide averages of pairwise genetic distances.

Although one potential explanation for increased divergence in low-recombination regions is that these regions are more resistant to post-divergence introgression (Brandvain et al. 2014), two observations are inconsistent with this explanation in *Neodiprion*: (1) D-statistics are not correlated with recombination rate (Fig. S4, Table S12), and (2) topological discordance is higher—not lower—in low-recombination regions. As expected, however, we did find a slight positive correlation between average D-statistic and Robinson-Foulds distance for the 500-kb window dataset (Fig. 6f), indicating that introgression does contribute to genome-wide patterns of discordance independent of other genomic features. A final piece of evidence that supports the hypothesis that positive selection contributes to heterogeneous gene-tree discordance across the genome is that average pairwise genetic distance—which should be maximized under divergent selection—was positively correlated with both gene density (Table S12; Fig. S4) and gene-tree discordance (Fig. 6d), regardless of window size (Table 1).

Together, our data suggest that gene-tree estimation error is both pervasive and contributes to heterogeneous discordance levels across the genome (Figs. 5 and 6, Table 1). To be clear, we are not claiming that gene-tree estimation error is the only source of discordance between gene trees and species trees in our dataset. For example, were the histories of each locus estimated with perfect accuracy, we would still expect considerable topological discordance due to incomplete lineage sorting and introgression. Rather, what our data indicate is that some regions of the genome (e.g., low-recombination regions with functionally important genes evolving under divergent selection) are more susceptible to gene-tree estimation error than others, elevating the level of discordance above and beyond that generated by incomplete lineage sorting and gene flow alone. And based on our data, this localized error is enough to outweigh localized reductions in effective population size that are expected to reduce topological discordance in low-recombination regions via reduced incomplete lineage sorting (Pease and Hahn 2013). However, because increased divergence in low-recombination regions (and therefore elevated gene-tree estimation error) is expected only when divergent selection is accompanied by gene flow, we hypothesize that a pattern of increased discordance in low-recombination regions may be unique to divergence-with-gene-flow scenarios.

Pervasive gene-tree estimation error also need not imply that the species-tree produced using those (erroneous) gene trees is wrong. For example, simulation studies indicate that summary methods that use gene trees as input can produce accurate species-tree estimates even when there is gene-tree estimation error (Larget et al. 2010; Liu et al. 2010a; Bayzid and Warnow 2013; Mirarab et al. 2014b, 2014a, 2016; but see Roch and Warnow 2015; Roch et al. 2019). Two observations from our own data also suggest that gene-tree estimation error likely did not have a large impact on our inferred species tree. First, species-tree estimates from ASTRAL were robust to window size (Fig. 2). Because different window partitioning strategies are likely to have very different sources of gene-tree estimation error—small windows lack information; large windows contain heterogeneous ancestries, base compositions, and substitution rates—robustness to window size suggests that these sources of error do not have a large impact on the inferred tree. Second, the SNP-based approach implemented in SVDquartets does not require gene-tree estimates at all, but produced a very similar species-tree estimate, albeit with minor differences in inferred relationships for three taxa that also had reduced quartet support in the ASTRAL tree (Figs. 2, 3). Still, it is possible that gene-tree estimation error—not sensitivity of methods to incomplete lineage sorting or introgression, as discussed above—is responsible for the observed differences between ASTRAL and SVDquartets trees.

Although our data reveal several genomic correlates that predict genome-wide and clade-wide patterns of discordance, there are several limitations of our dataset that stem from using a single reference genome and clade-wide metrics for genomic variables that may vary among taxa or groups of taxa. For example, variation in gene position and copy number among species could have introduced error into our estimates of local gene density, recombination rate, and GC content, all of which were based on the *N. lecontei* reference genome. Similarly, we estimated local recombination rates from a single inter-population cross in *N. lecontei* (Linnen et al. 2018). Although recombination rate is conserved among some closely related species, it is rapidly evolving in others (Smukowski and Noor 2011). It would therefore be worthwhile to generate additional reference genomes and estimate recombination rates in additional populations and species to further evaluate the robustness of our conclusions. We also used clade-wide averages of pairwise genetic distances and D-statistics, and a tree-wide measure of topological discordance. However, patterns and correlates of gene-tree discordance could vary among different parts of the tree. For example, the impact of introgression on topological discordance should be most pronounced in lineages that experienced gene flow. Analyzing subsets of the tree and collecting additional data to better characterize genomic predictors of discordance (e.g., introgression analyses with multiple individuals per species; recombination rate estimates for additional species) would be fruitful avenues for future research. Despite these limitations, our results demonstrate how a multiple regression approach can be used to better understand heterogeneous phylogenetic signal. Similar analyses in other taxa may reveal whether sources of discordance vary predictably among taxa with different divergence histories. Essential ingredients for these analyses are high-quality, annotated reference genomes, which are increasingly available for non-model organisms (Hotaling et al. 2021).

### A Discordance Hotspot on Chromosome 1

Although we have identified several genomic predictors of topological discordance, these do not appear to explain a ∼3.5 Mb discordance hotspot on Chromosome 1 (Fig. 5). For example, although Robinson-Foulds distances calculated for the seven 500 kb windows located in the discordance hotspot are much higher than all but one other 500 kb window in the genome, these windows do not stand out in terms of any of the genomic predictor variables we evaluated (yellow points, Fig. 6). That said, while the discordance hotspot does not have unusual recombination rates as inferred from a mapping cross in *N. lecontei* (Fig. 6e), the hotspot is both unusually large and robust to window size (Fig. 5). By comparison, other regions with high Robinson-Foulds distances tended to appear only when window sizes were small and were primarily in data-poor centromeres or telomeres (Figs. 5c, 5d). Based on these observations, we hypothesize that the discordance hotspot may correspond to the location of a genomic rearrangement segregating within or between species (but not the mapping population used to estimate recombination rates) that reduces recombination in this region, thereby leaving a large footprint of discordant signal. Evaluating this hypothesis will require characterizing genomic rearrangements across the genus, which will require *de novo* assemblies from additional populations and species rather than anchoring analyses to a representative assembly.

Although mechanisms that reduce recombination can explain why windows sampled across a reasonably large region of a chromosome can produce similar topologies (Figs. 5, 7b), reduced recombination does not explain why that topology is so different from the underlying species tree. One possibility is that this location is a hotspot for adaptive introgression. Consistent with this explanation, average genome-wide D-statistics estimated for this region tended to be on the higher-end compared to the rest of the genome (quantile-normalized values > 0), but there are also less-discordant regions with even higher introgression estimates (Fig. 6f). A non-mutually exclusive explanation for unusually strong discordance in this region is that it is a hotspot for adaptation via ancestral polymorphism. Determining the relative contribution of introgression and incomplete lineage sorting to discordance in this region will require additional sampling and analysis to characterize the history of divergence, selection, and gene flow among *Lecontei* group species and across their genomes.

As a starting point to generate more specific adaptive hypotheses for the discordance hotspot, we mapped two ecologically important traits that are variable across the *Lecontei* clade onto the consensus species tree (Fig. 7a). Although we have not reconstructed ancestral character states, the distribution of characters across the tree cannot be explained without multiple independent transitions between different diapause strategies (egg, prepupa, or adult overwintering) and independent gains or losses of different pine hosts. Thus, at least some shared diapause traits and host associations must be due to convergent phenotypic evolution. Shared diapause strategies and hosts are also likely to result in convergent selection pressures due to shared climatic challenges, host characteristics, and predator/parasite communities, as well as more opportunities for hybridization due to increased temporal and spatial overlap. If the discordance hotspot is in some way related to convergent diapause or host-use strategies—either as the genetic source of these convergent traits or as a target of correlated selection pressures— we would expect the species to cluster by trait in the discordance-hotspot tree. This prediction is partially supported. For example, all species that do not overwinter as prepupae in cocoons cluster together in a single clade in the consensus tree for the seven 500 kb windows located in the discordance hotspot (*pinusrigidae* complex + *pratti* complex + *N. warreni*; Fig. 7b). There is also some clustering by host use in this tree, with jack pine-specialist *N. rugifrons* falling into a clade containing three other species that also use jack pine (*N. abbotii, N. dubiosus, N. nigroscutum*).

Gene annotations may provide additional clues about the nature of the discordance hotspot. Based on gene names alone, we identified at least eight genes involved in chemosensation in the high-discordance region that could be important for host use, as well as six genes that could play a role in premating and postcopulatory interactions (Table S13). Intriguingly, this region also contained at least nine genes that could be involved in mitochondrial-nuclear interactions (Table S13). Given a history of pervasive mitochondrial introgression in *Neodiprion*, we compared our discordance-hotspot consensus tree (Fig. 7b) with a mtDNA consensus tree estimated from a 1752 bp alignment spanning the mitochondrial genes *cytochrome c oxidase I, tRNA-leucine*, and *cytochrome c oxidase II* sequenced in 123 individuals (1-14 populations from each of the 19 *Lecontei* group species; Linnen and Farrell 2007) (Fig. 7c). Although the two trees were not identical, there were some striking similarities, including an unusual clade containing *N. warreni*, all *pratti* complex members, and two *pinusrigidae* complex members, and a clade containing a mix of *virginianus* and *abbotii* complex members (Fig. 7b vs. Fig. 7c). One potential explanation for this combination of observations—specifically, high rates of mitochondrial introgression, similarity of the mtDNA tree and the discordance-hotspot tree, and gene annotations consistent with mito-nuclear interactions—is that the discordance hotspot contains genes involved in mito-nuclear incompatibilities, which are a common source of intrinsic postzygotic isolation (Burton and Barreto 2012; Sloan et al. 2017). Under this explanation, introgression of a particular mitochondrial haplotype would require co-introgression of compatible alleles at interacting nuclear loci, yielding similar gene-tree topologies for the co-introgressed regions. Although additional data are needed to test this model, such data would support the hypothesis that gene trees for speciation genes—specifically, loci involved in Dobzhansky-Muller incompatibilities—are less likely to be concordant with the species tree than other loci (Wang and Hahn 2018).

### Conclusions

Over 15 years ago, we documented pervasive mitochondrial introgression in *Neodiprion* involving many members of the eastern *Lecontei* clade, rendering mitochondrial data unreliable for species-tree inference (Linnen and Farrell 2007). A “multi-locus” dataset of three nuclear genes was, unsurprisingly, insufficient for generating a robust species-tree estimate (Linnen and Farrell 2008a). Here, a whole-genome alignment analyzed with contemporary methods has produced a well-resolved species-tree that—except for three uncertain relationships—is robust to locus-sampling and analysis strategy. Unexpectedly, patterns of topological discordance across the nuclear genome, which we attribute partly to local genomic features that increase gene-tree estimation error, revealed a striking window of gene-tree discordance on Chromosome 1. This region contains several genes that are likely to interact with mitochondrial genes, and gene trees estimated from the discordance hotspot resemble the mtDNA tree. Thus, we have come full circle back to mitochondrial introgression, now with possible co-introgression of interacting nuclear loci in the discordance hotspot, which we hypothesize may be locked together in an inversion that reduces recombination. Overall, these findings demonstrate the utility of combining phylogenomic analysis with high-quality reference genomes and thorough data exploration for generating novel hypotheses about the molecular mechanisms and evolutionary processes that produce variable genealogical histories across genomes.

## Acknowledgements

We thank members of the Linnen and Weisrock labs for helpful discussion. This work was supported by the University of Kentucky Center for Computational Sciences and the Lipscomb High Performance Computing Cluster, the United States Department of Agriculture National Institute of Food and Agriculture (2016-67014-2475; CRL), the National Science Foundation (DEB-CAREER-1750946; CRL), and a University of Kentucky Lyman T. Johnson Fellowship (KV).

## Data Availability Statement

The *Neodiprion lecontei* reference genome assembly and annotation (and associated reads) are available on NCBI (NCBI BioProject PRJNA784628, accession number GCA_021901455.1). For remaining species, all sequencing reads will be available on NCBI upon publication (NCBI BioProject PRJNA854171, accession numbers SRR19909288-SRR19909306). *Neodiprion* pseudo-reference genomes (aligned genomes for each species in *Neodiprion lecontei* genome coordinates) will be available on Dryad. Custom scripts for generating *Neodiprion* pseudo-reference genomes, converting reference and pseudo-reference genomes into datasets for analysis (windows, genes, and SNPs), and all downstream analyses will be available on the LinnenLab GitHub page (https://github.com/LinnenLab) under the Herrig_etal_NeodiprionPhylogeny repository.

