## Supplemental Figures for "Whole Genomes Reveal Evolutionary Relationships and Mechanisms Underlying Gene-Tree Discordance in *Neodiprion* Sawflies"

Map Reads to

*N. lecontei* genome

(bowtie2)

Incorporate Invariant Sites

(samtools and bcf tools)

Map to current iteration of species genome

Incorporate Invariant Sites

(samtools and bcf tools)

Repeat 4 times

Determine Depth at Each Site and maximum cutoffs

Map to current iteration of species genome

Incorporate “N”s for sites with a depth below 4 or greater than the maximum

Final Pseudo-Reference Based Genome

**Figure S1.** Pipeline for pseudo-reference genomes produced using the *N. lecontei* reference genome and short-read data from 19 additional species.

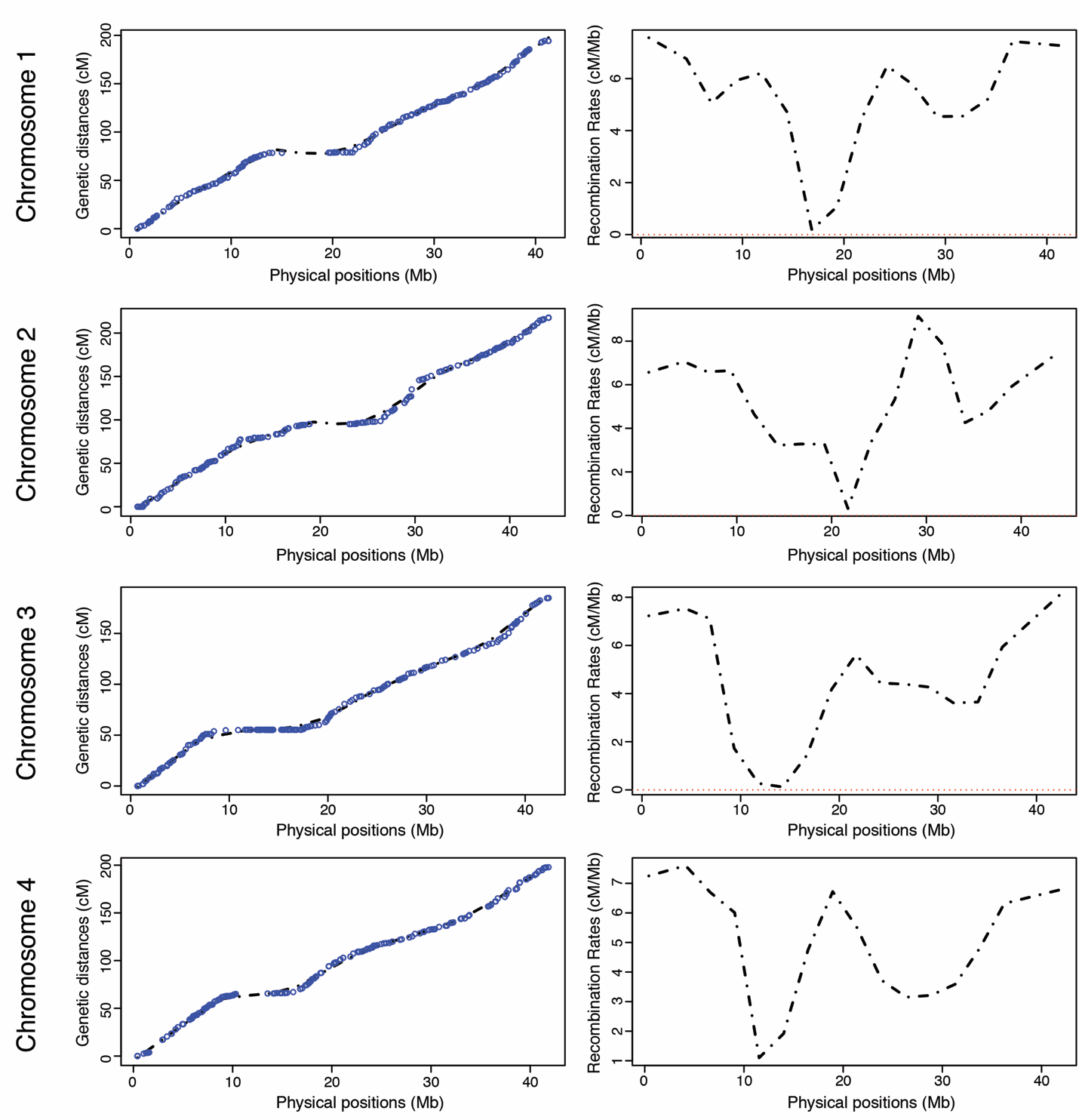

**Figure S2:** Genetic map and local recombination rate for chromosomes 1-4. Panels on the left depict the relationship between physical distance and genetic distance, with individual markers shown as blue dots and model fitted via sliding windows (dotted line). Panels on right depict the recombination rate (cM/Mb) as a function of physical distance.

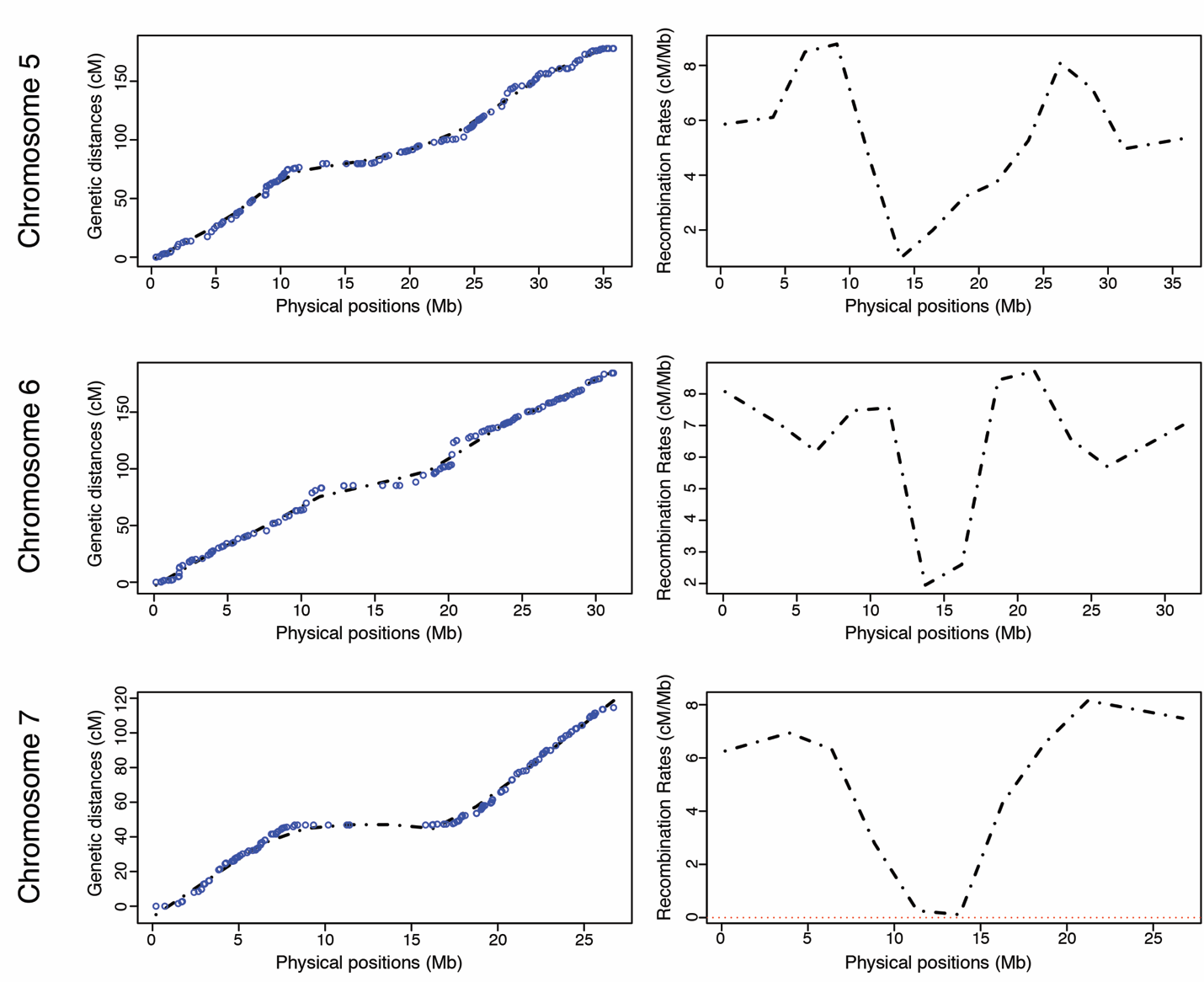

**Figure S3:** Genetic map and local recombination rate for chromosomes 5-7. Panels on the left depict the relationship between physical distance and genetic distance, with individual markers shown as blue dots and model fitted via sliding windows (dotted line). Panels on right depict the recombination rate (cM/Mb) as a function of physical distance.

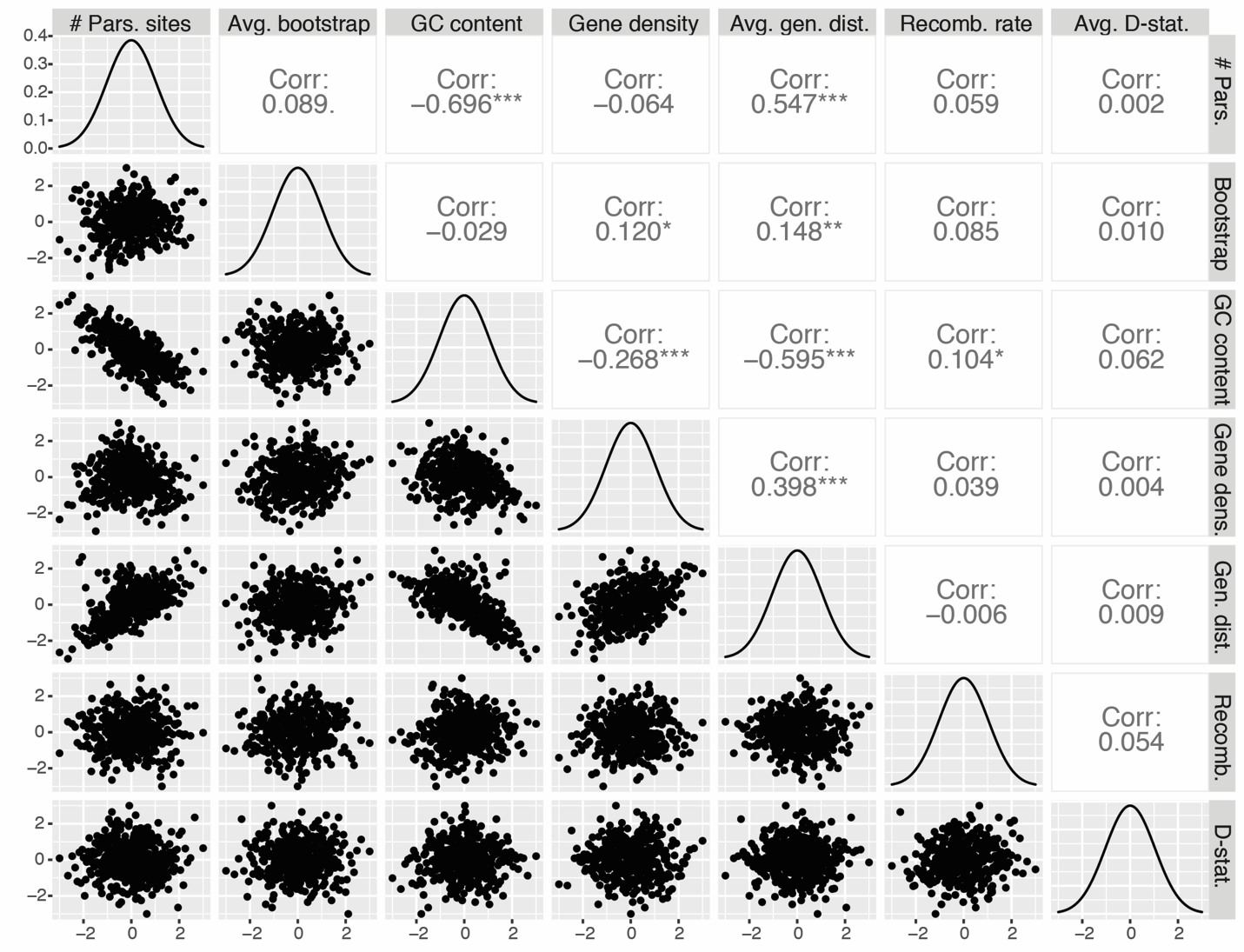

**Figure S4:** Correlation matrix for genomic predictors of discordance estimated in 500 kb windows. From left-to-right and top-to-bottom, predictors are: number of parsimony-informative sites, average gene-tree bootstrap support, GC content, gene density, average pairwise genetic distance (K80 distance), average D-statistic (ABBA-BABA), recombination rate (cM/MB). All values are normal-quantile transformed to better visualize relationships between variables. Below the diagonal: pairwise scatter plots; above the diagonal: Pearson’s correlation coefficient and significance (****P <* 0.001; ***P*<0.01; **P<*0.05; *.P*<0.10)*;* on the diagonal: distributions for variables.
